# A human induced pluripotent stem (hiPS) cell model for the holistic study of epithelial to mesenchymal transitions (EMTs)

**DOI:** 10.1101/2024.08.16.608353

**Authors:** Caroline Hookway, Antoine Borensztejn, Leigh K. Harris, Sara Carlson, Gokhan Dalgin, Suraj Mishra, Nivedita Nivedita, Ellen M. Adams, Tiffany Barszczewski, Julie C. Dixon, Jacqueline H. Edmonds, Erik A. Ehlers, Alexandra J. Ferrante, Margaret A. Fuqua, Philip Garrison, Janani Gopalan, Benjamin W. Gregor, Maxwell J. Hedayati, Kyle N. Klein, Chantelle L. Leveille, Sean L. Meharry, Haley S. Morris, Gouthamrajan Nadarajan, Sandra A. Oluoch, Serge E. Parent, Amber Phan, Brock Roberts, Emmanuel E. Sanchez, M. Filip Sluzewski, Lev S. Snyder, Derek J. Thirstrup, John Paul Thottam, Julia R. Torvi, Gaea Turman, Matheus P. Viana, Lyndsay Wilhelm, Chamari S. Wijesooriya, Jie Yao, Julie A. Theriot, Susanne M. Rafelski, Ruwanthi N. Gunawardane

## Abstract

The epithelial to mesenchymal transition (EMT) is a widely studied but poorly defined state change due to the variety of ways in which it has been characterized in cells. There is a need for reproducible cell model systems that enable the integration and comparison of different types of measured observations of cells across many distinct cellular contexts. We present human induced pluripotent stem (hiPS) cells as such a model system by demonstrating its utility through a comparative analysis of hiPS cell-EMT in 2D and 3D cell culture geometries. We developed live-imaging-based assays to directly compare examples of changes in cell function (via migration timing), molecular components (via expression of marker proteins), organization (via reorganization of cell junctions), and environment (via dynamics of basement membrane) in the same experimental system. The EMT-related changes we measured occurred earlier in 2D colonies than in 3D lumenoids, likely due to differences in the basement membrane environments associated with 2D vs. 3D initial hiPS cell culture geometries. We have made the 449 60-hour-long 3D time-lapse movies and the associated tools used for analysis and visualization open-source and easily accessible as a resource for future work in this field.

## Introduction

Cell state transitions are fundamental to biological processes, but because cell states can be described by a variety of cell properties, cell states and cell state transitions are very difficult to define^1^. Defining the epithelial to mesenchymal transition (EMT), a state change that occurs in several biological contexts from embryonic development to cancer metastasis, is an example of this challenge^2–4^. For instance, EMT can be described *functionally* as the loss of the barrier-forming ability of cells in the epithelial sheet and the gain in migratory and invasive behavior of mesenchymal cells^5,6^. EMT can be described *organizationally* as a change in the apicobasal polarity of epithelial cells to a gain in the front-rear organization of a migrating cell, including changes in cell adhesion and cell-cell junctions^7,8^. EMT can be described as changes in the *molecular census* of the cell comprising changes in protein and transcription factor expression, including, for example, expression of T-BOX transcription factors^3,9–11^. Finally, EMT can be described by changes in how cells interact with their *environment*, both with the extracellular matrix (ECM) and each other. For example, the basement membrane, a specialized sheet of ECM that underlies most animal tissues, is penetrated by migrating mesenchymal cells as they migrate to a new environment^12^. However, in individual experimental studies, EMT is often defined within only one of these types of descriptions, and there is little knowledge of the relationships between functional, organizational, and molecular census observations over the course of EMT.

The goal of this study was to establish a controlled system in which to study the interactions between different types of observations that together define the holistic cell state change that is EMT. Human induced pluripotent stem (hiPS) cells serve as a powerful model for this exploration because they are a normal human cell type that grow in culture without immortalization or karyotypic abnormalities, and the differentiation of hiPS cells provides a unique window into the orchestrated dynamics of EMT observed during early embryonic development^13^. EMT can be induced in hiPS cells by activating the Wnt pathway with growth factors or small molecules^14,15^, resulting in an EMT that resembles gastrulation following primitive streak formation^16^. Additionally, hiPS cells can be cultured in multiple different geometries, including 2D colonies made of epithelial-like sheets, and 3D lumenoids that are hollow spheres made up of a monolayer of epithelial-like cells oriented with their apical faces contacting a central lumen and their basal sides pointing outward^17,18^. 2D colonies can be converted to adherent 3D lumenoids with the application of dilute mouse-derived basement membrane proteins in the form of Matrigel to the surrounding media^19^. The ability to culture hiPS cells in different starting cell culture geometries provides an experimentally standardized model in which to study EMT, enabling data collection across multiple types of observations within the same cell model as well as the comparison of EMT across multiple geometrical contexts. This kind of simplified system is needed because understanding the EMT that occurs during gastrulation in an animal embryo is complicated by the developmental patterning that serves as the overwhelming arbiter of many cell decisions, behaviors, and fate transitions, including mechanical stress and signaling gradients^20,21^.

We used the timing of cell migration, expression of EMT-related and stemness-related transcription factors, and the reorganization and loss of cell-cell junctions as representative readouts of changes in function, molecular census, and cell organization respectively, during EMT from 2D and 3D hiPS cell culture geometries. Except for the loss of stemness marker SOX-2, we found that the timing of these changes occurred earlier in 2D conditions compared to 3D, revealing critical insights into how the dimensionality of the cell culture system influenced EMT. Importantly, we were also able to investigate the relationship between these changes and the basement membrane. Using a live-imaging approach to visualize hiPS cell-derived collagen IV during EMT, we found that hiPS cells cultured in 3D produce and surround themselves with a basement membrane shell that cells must cross in order to migrate away. We found that the presence of this basement membrane appears to cause a delay in the observed time of migration by blocking egress of cells that have already delaminated, an effect we do not observe in 2D conditions where the basement membrane is much less established. These results demonstrate the utility of this hiPS cell-based model for studying EMT in a holistic way.

## Results

### An hiPS cell-based model for comparing EMT in 2D and 3D environmental contexts

We developed a chemically-induced EMT in human induced pluripotent stem (hiPS) cells as a model system to follow multiple distinct types of observations during the EMT state change from hiPS cells grown in different starting cell culture geometries. To demonstrate the utility of this model, we generated a streamlined and reproducible live-cell imaging and analysis assay. We started with an hiPS cell line with mEGFP-tagged histone H2B to visualize nuclei along with a fluorescent dye-conjugated human-specific anti-collagen IV antibody^22^ to visualize the basement membrane specifically produced by the hiPS cells, and not the mouse-derived basement membrane material (Matrigel) present in the system. This allowed us to observe the re-organization of cells, changes in the basement membrane, and cell-basement membrane interactions throughout the EMT process.

The 2D condition consisted of hiPS cells cultured on Matrigel-coated glass (Fig. 1 row 1 *hiPS cell 2D colony growth*). In this condition, cells grew as epithelial-like 2D monolayer colonies with their basal sides oriented towards the glass. These colonies expanded as cells proliferated and produced very little collagen IV (Fig. 2A and C top row). We created 3D hiPS cell culture geometries by exposing 2D colonies on Matrigel-coated glass to dilute Matrigel in the surrounding medium leading to the formation of 3D lumenoids (Fig. 1 row 5, *2D to 3D lumenoid formation*). Time-lapse imaging of the transition from 2D colony to 3D lumenoid in the presence of the human-specific anti-collagen IV antibody revealed the role of the basement membrane in this process: Anti-collagen IV labeling was first observed at colony edges, suggesting that edge cells were lifting their basal sides up and away from the glass surface. After lifting off the glass, the cells at the edges of the colony then cinched closed in a purse string-like mechanism, eventually closing completely to form a single central lumen ∼24 h after Matrigel was introduced (Fig. 2B and C bottom row). As cells continued to proliferate and grow, the lumen appeared to inflate and the whole structure became spherical. In contrast to colony growth, where little collagen IV labeling was observed, a bright, thick shell of collagen IV surrounded the lumenoid, demonstrating that lumenoid cells deposit basement membrane as they grow (Fig. 2B). This transformation from 2D colony to 3D lumenoid was very robust and reproducible, consistently resulting in lumenoids that were spherical yet firmly anchored to the glass, facilitating downstream media changes and long-term time-lapse imaging in 96-well format.

**Fig. 1:**
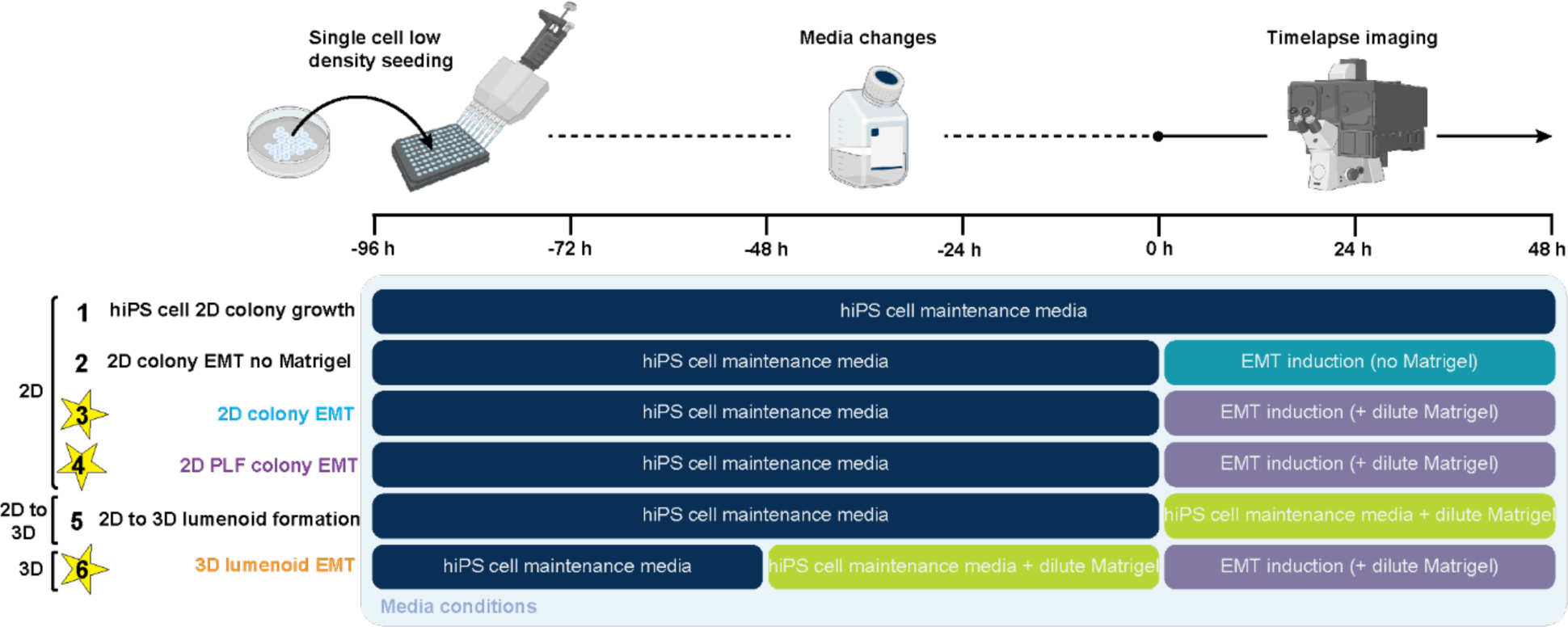
Overview of hiPS cell growth and EMT conditions for imaging assay. Schematic showing hiPS cell growth conditions in 2D cell culture geometries (row 1), the formation of 3D lumenoids (row 5), and EMT induction conditions in 2D (rows 2, 3 and 4) and 3D (row 6) cell culture geometries. The EMT conditions compared throughout the remainder of the manuscript are starred. See Methods section 1 for specific details on cell seeding and media preparation.

**Fig. 2:**
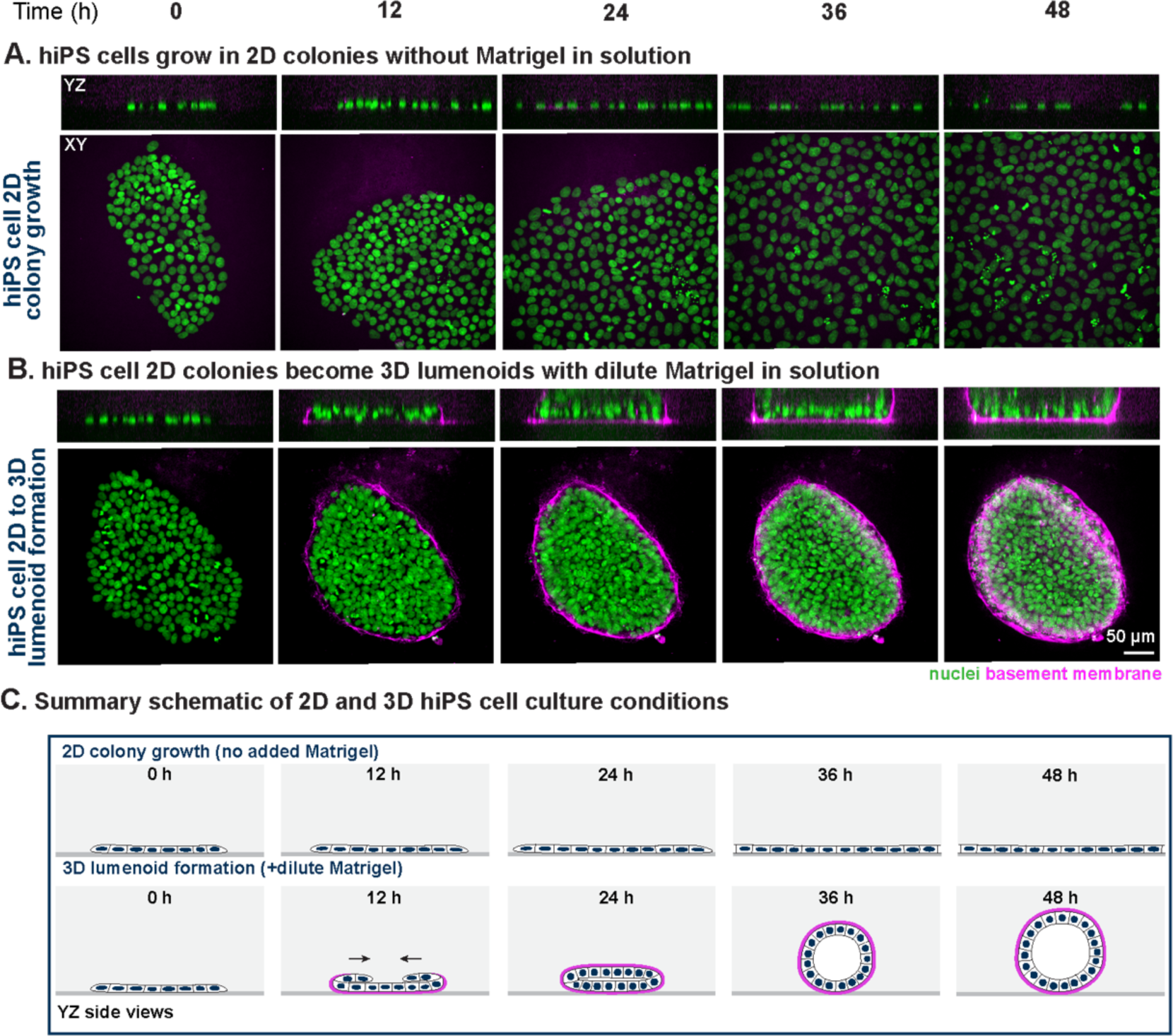
Environmental Matrigel changes hiPS cell colony geometry during growth. **A.** Representative example frames at 12 h intervals of *hiPS cell 2D colony growth* from 30-minute interval time-lapses showing flat expansion of the colony as seen by the nuclei (mEGFP-tagged H2B; green) with little hiPS cell-derived basement membrane (human-specific anti-collagen IV antibody; magenta). XY views show maximum intensity Z-projections. YZ views are single Z-slices through the approximate middle. **B.** Same as A for *2D to 3D lumenoid formation* showing transition from a flat epithelial sheet to a 3D lumenoid surrounded by hiPS cell-derived basement membrane when grown in culture medium containing dilute Matrigel (see Methods section 1.3). Scale bar: 50 µm. **C.** Schematic representation of *hiPS cell 2D colony growth* and *2D to 3D lumenoid formation* with nuclei as filled navy ellipses, cell boundaries as black lines, and basement membrane as magenta lines. See Supplemental Table 1 for web-based 3D Volume Viewer links for each time-lapse shown.

Next, we induced EMT in these 2D and 3D hiPS cell cultures by downstream activation of the Wnt pathway using small molecule (CHIR99021) inhibition of GSK3β (Methods section 1.4). In 2D colonies grown on Matrigel-coated glass (Fig. 1 row 2, *2D colony EMT no Matrigel*), cells contracted and heightened until ∼20-30 h, after which they migrated away from the initial colony (Fig. 3A). Collagen IV labeling was seen only at the glass surface (Fig. 3A). However, we excluded this *2D colony EMT no Matrigel* condition from the rest of the study due to frequent cell death. In contrast, when dilute Matrigel was included in the induction medium (Fig. 1 row 3, *2D colony EMT*), there was consistently less cell death. In this condition, collagen IV labeling was seen at the colony periphery and a fully-formed but not yet spherical lumenoid was formed within the first ∼24 h post EMT induction, consistent with dilute Matrigel driving lumenoid formation (in the presence or absence of EMT induction). This nascent lumenoid was only weakly coated with collagen IV before cells migrated out and away at ∼20-30 h (Fig. 3B).

**Fig. 3:**
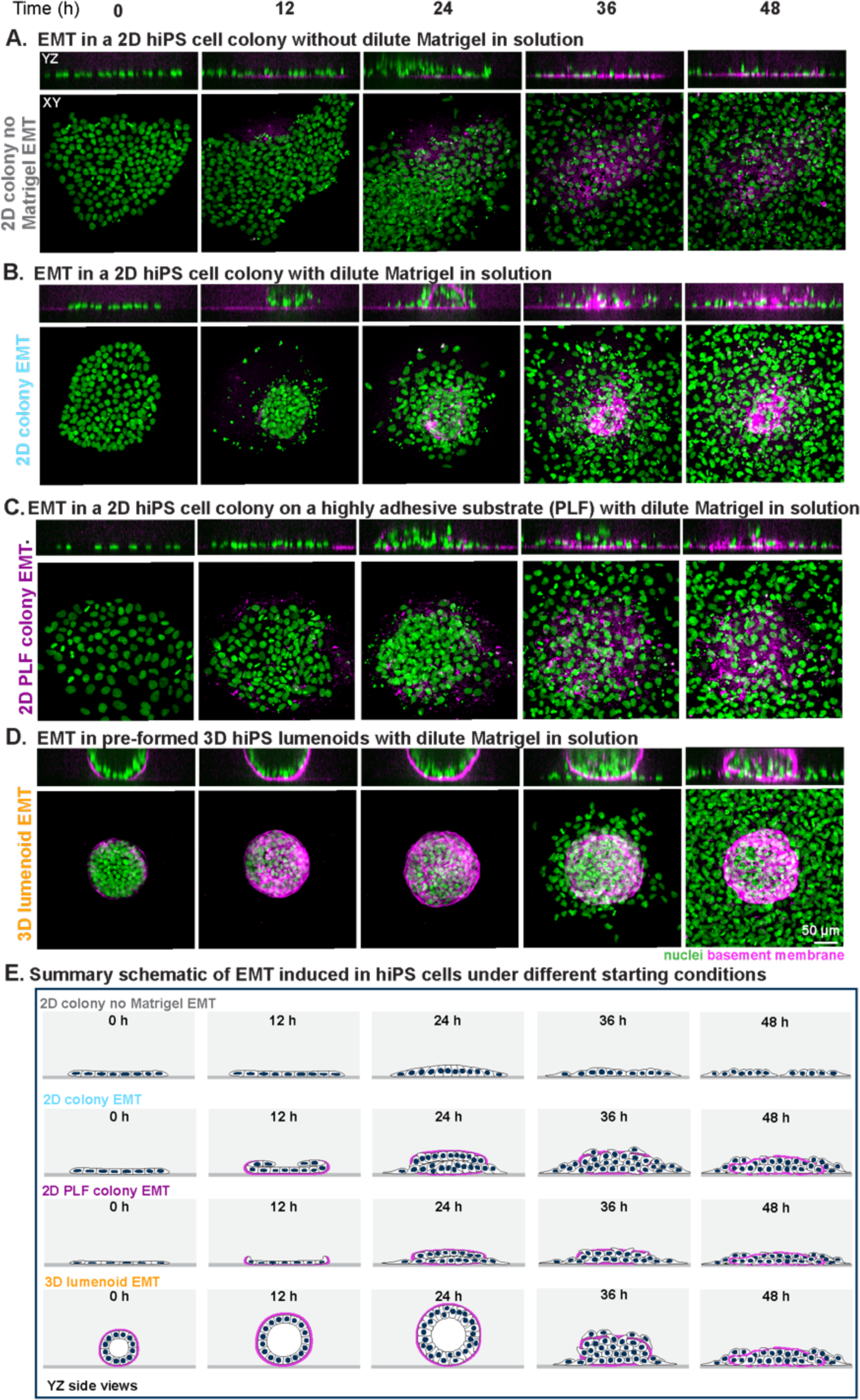
EMT is influenced by starting hiPS cell colony geometry, environmental Matrigel, and substrate adhesiveness. In sample images, nuclei are marked by mEGFP-tagged H2B (green) and the basement membrane (human-specific anti-collagen IV antibody) is visualized in magenta. **A.** Upon EMT induction, cells from 2D hiPS cell colonies plated on Matrigel substrate without Matrigel in solution (*colony EMT no Matrigel,* selected healthy example) disperse (as seen by nuclei) and migrate away, remaining flat and depositing little hiPS cell-derived basement membrane. XY views show maximum intensity Z-projections. YZ views are single Z-slices through the approximate middle. **B.** hiPS cells grown on Matrigel with simultaneous introduction of dilute Matrigel to the solution during EMT induction (*colony EMT*) increase basement membrane deposition and begin forming a lumenoid, which collapses upon migration onset. **C.** hiPS cells plated on PLF instead of Matrigel with simultaneous dilute Matrigel addition and EMT induction (*2D PLF colony EMT*) are flatter and more spread out, forming flatter lumenoids prior to migration. **D.** Cells from pre-formed 3D lumenoids plated on Matrigel substrate (*3D lumenoid EMT*) migrate through the basement membrane and away from the collapsing lumenoid. Scale bar: 50 µm. **E.** Schematic outlining colony dynamics during EMT for different starting colony conditions. Schematics are drawn as YZ side views with nuclei represented as filled navy ellipses, cell boundaries as black lines, and basement membrane as magenta lines. See Supplemental Table 1 for web-based 3D Volume Viewer links for each time-lapse shown.

We also induced EMT in the presence of dilute Matrigel from colonies plated on a different substrate, poly-D-lysine, laminin, and fibronectin (PLF) (Fig. 1 row 4 *2D PLF colony EMT*). Cells in colonies grown on PLF appeared flatter and more spread than those on Matrigel, suggesting that PLF was the more adhesive of the two substrates^23^ (Fig. 3 0 h panels in A and B vs. 0 h panel in C). We called this condition *2D PLF colony EMT*. *2D PLF colony EMT* looked similar to *2D colony EMT*, including the timing of initial lumenoid formation and cell migration (Fig. 3C. vs. B.). However, the nascent lumenoid remained very flat in *2D PLF colony EMT* (Fig. 3C 36 and 48 h panels).

Finally, we also induced EMT in pre-formed lumenoids (Fig. 1 row 6 *3D lumenoid EMT*). After induction, lumenoids continued to grow as hollow spheres for ∼24 h (Fig. 3D 0 to 24 h panels), after which the lumen disappeared, and cells migrated across the basement membrane and out onto the glass. The basement membrane surrounding the pre-formed lumenoid in *3D lumenoid EMT* was brighter and thicker than the shell that surrounded the nascent lumenoid formed in *2D colony EMT* or *2D PLF colony EMT* (Fig. 3D vs. 24 h panel of B and C). The two-day exposure of colonies to dilute Matrigel to pre-form the lumenoids in *3D lumenoid EMT* (Fig. 1 row 6) likely provided more time for cells to deposit basement membrane prior to EMT induction than in the 2D conditions.

Thus, by manipulating the adhesiveness of the substrate and the environmental Matrigel, we were able to grow hiPS cells in different colony geometries: flat 2D colonies on a Matrigel substrate (*2D colony EMT*), even flatter 2D colonies on a PLF substrate (*2D PLF colony EMT*), and round, 3D lumenoids on a Matrigel substrate with dilute Matrigel in solution (*3D lumenoid EMT*). These provided a set of three different starting points from which to compare EMT, allowing us to create the *2D colony EMT*, *2D PLF colony EMT*, and *3D lumenoid EMT* conditions which are compared in the remainder of this study (Fig. 1, starred rows 3, 4, and 6; Fig. 3B, C and D).

The hiPS cell-EMT model system together with this image-based assay and these specific conditions permitted us to create a high-quality, standardized, and well-annotated dataset for downstream analysis. This dataset consists of 449 individual movies systematically acquired via live-cell time-lapse imaging every 30 min for 60 h over six cell lines (Methods section 2.2). Time-lapses were selected for analysis based on criteria of cell health and the lack of migrating cells from secondary colonies or lumenoids and only the first 48 h of imaging were included in analysis, resulting in a set of 268 48-hour analyzed time-lapses (Methods section 3.2).

### Live-cell visualization of migrating cells crossing the basement membrane reveals cell-basement membrane interactions

The ability to observe cells and basement membrane together in live-cell time-lapse images of EMT from 2D and 3D hiPS cell culture geometries made it possible to investigate the relationship between cells and their ECM environment. Cell and basement membrane interactions were more visible in the basement membrane of the 3D lumenoid because it was brighter and thicker than in either of the 2D conditions, presumably because it had more time to be formed. This permitted a deeper visual analysis of the interactions between the basement membrane and the cells during *3D lumenoid EMT*. An AGAVE 3D rendering of the nuclei and basement membrane during *3D lumenoid EMT* shows the well-formed basement membrane at 0 h, the wavy deformations in the basement membrane wall prior to cell migration at 24 h, the migration of cells through the basement membrane by 36 h, and the “shell” of basement membrane left behind once the majority of cells have become migratory by 48 h (Fig. 4A). A maximum intensity projection of the basement membrane shell reveals the formation and enlargement of holes (Fig. 4B). Inspection of single nuclei in the raw images reveals shape changes in initial cells that exit through the basement membrane, narrowing as though they were squeezed through (Fig. 4C and Supplemental Movie 1). Later, we saw the appearance of larger holes in the basement membrane and less change in shape of nuclei that crossed through these holes (Fig. 4D and Supplemental Movie 2). Interestingly, we saw that after the initial exit, cells occasionally crossed through holes in the basement membrane in both outwards and inwards directions (Supplemental Fig. S1 and Supplemental Movie 3). Since exit through the basement membrane was difficult to compare across 2D and 3D conditions due to differences in basement membrane thickness and morphology, we next turned to other ways to characterize EMT across conditions.

**Fig. 4:**
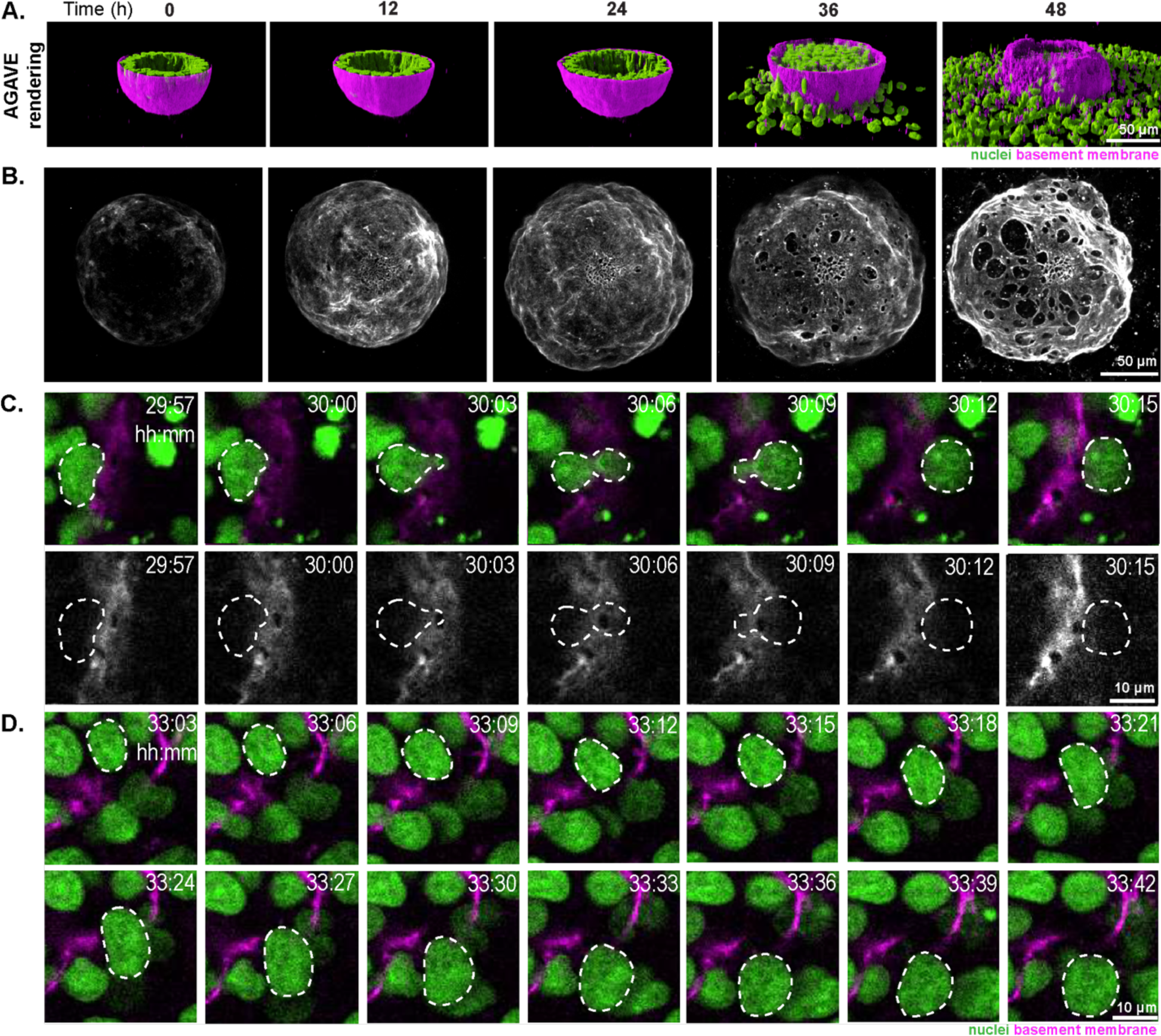
Cells can leave lumenoids by deforming to fit through narrow holes in the basement membrane or by passing, undeformed, through larger holes. In all 3D renderings and colored sample images, nuclei are marked by mEGFP-tagged H2B (green) and the basement membrane (human-specific anti-collagen IV antibody) is visualized in magenta or white. **A.** AGAVE 3D rendering of images in Fig. 3D, showing cells (nuclei; green) leaving the lumenoid through the basement membrane (magenta). **B.** Maximum intensity Z-projection of basement membrane (gray) from the example shown in magenta in Fig. 3D and 4A. All images (C-D) are single Z-slices at the approximate middle, shown in XY. **C.** Three-minute time-lapse imaging of *3D lumenoid EMT* showing a nucleus deforming as though it was squeezed through a small opening in the basement membrane. A dashed line outlines the deforming nucleus of interest. The basement membrane channel of the example in the top row is shown below in gray. Note the hole in the basement membrane at the narrowest part of the nucleus as it exits the lumenoid. Timestamps: time post-EMT induction (see Supplemental Movie 1). **D.** An example similar to C where a cell is seen exiting through a larger opening in the basement membrane of a 3D lumenoid with minimal nuclear deformation. Scale bars: 10 µm (see Supplemental Movie 2). See Supplemental Table 1 for web-based 3D Volume Viewer links for each time-lapse shown.

### Cells migrate earlier in the 2D EMT conditions vs. *3D lumenoid EMT* in hiPS cells

We compared the timing of cell migration between the two 2D EMT conditions and *3D lumenoid EMT* to ask whether differences in starting colony geometry could influence when cells begin to migrate. First, we took advantage of the bright-field channel included in all of the data in the dataset across the six fluorescently tagged cell lines and developed an image analysis approach to create a 3D segmentation mask demarcating the combined total volume of all of the cells together (vs. background) in each of the volume stacks across the dataset (Fig. 5A). To generate these “all-cells masks,” we trained a deep learning model with 3D image stacks of ground truth masks generated from cytoplasmic mEGFP-expressing cells paired with their respective 3D bright-field images (Methods section 4.2, Supplemental Fig. S2). Next, because migrating cells spread out onto the glass as they migrated, we could measure changes in the area of the combined cells’ footprint at the glass surface using a maximum intensity Z-projection of the bottom two Z-slices of each all-cells mask (Fig. 5B).

**Fig. 5:**
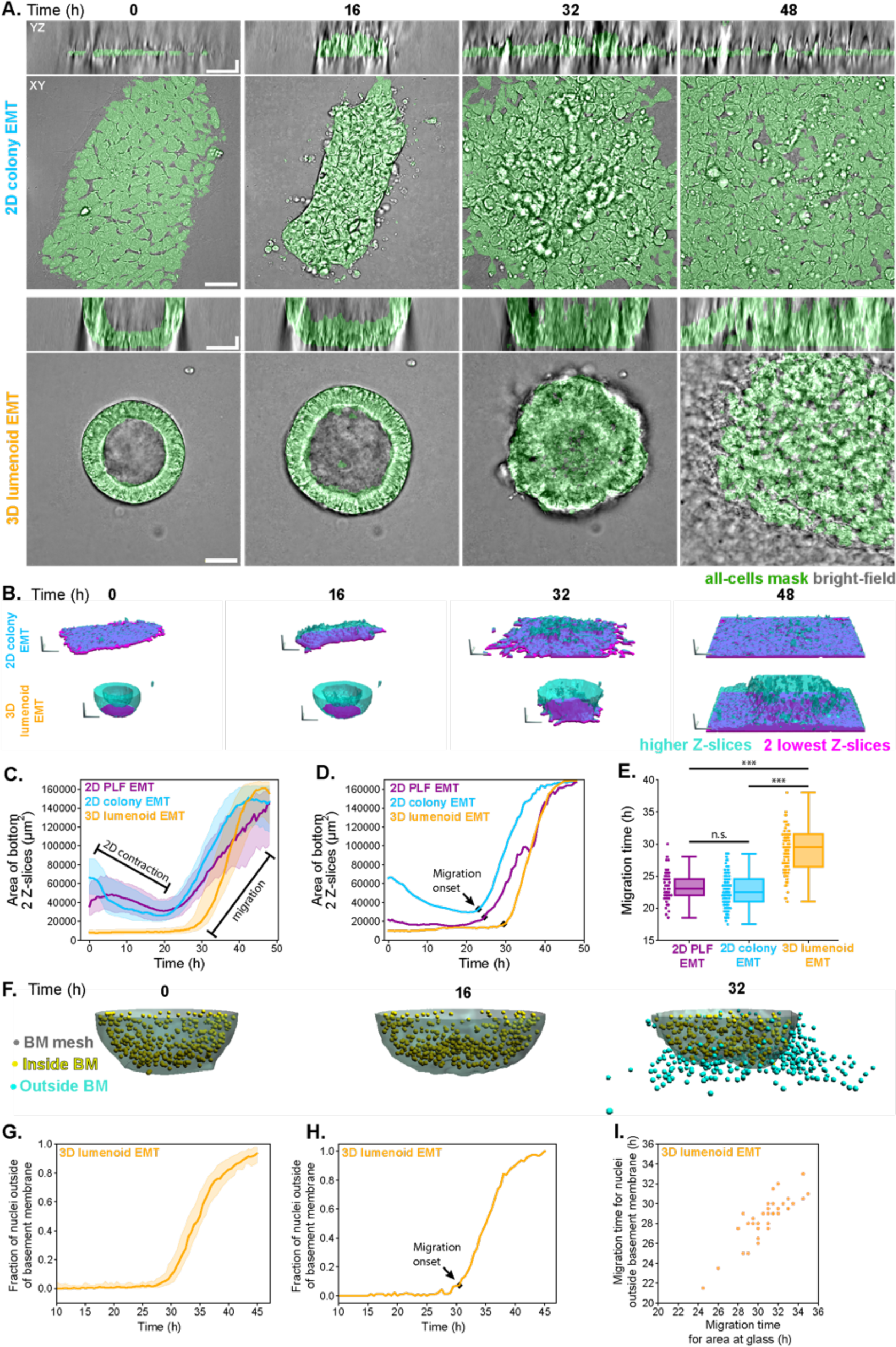
Migration timing differs for 2D colony and 3D lumenoid geometries as determined by all-cells mask-derived systematic measurements of colony spreading. **A.** Overlay of transmitted light (grey) and the corresponding all-cells masks (green) at a single Z or X slice for *2D colony EMT* (top) and *3D lumenoid EMT* (bottom) conditions. Scale bars: 50 µm (horizontal) and 20 µm (vertical). **B.** 3D renderings of bright-field-generated masks with the bottom two Z-slices shown in magenta and all others shown in translucent cyan. Note that the model successfully predicts and segments the hollow nature of the *3D lumenoid EMT* condition. Scale bars: 50 µm per axis. See Supplemental Table 1 for web-based 3D Volume Viewer links for each time-lapse shown. **C.** Area of the maximum intensity projection of the bottom two Z-slices of all-cells masks (i.e. colony area at the glass; magenta in B) over time. The line is the median and the shaded region is the 25^th^-75^th^ percentile range for the *Colony EMT* (n=63), *2D PLF colony EMT* (n=59), and *3D lumenoid EMT* (n=83). **D.** Example of individual trajectories from C of colony area at the glass over time showing computed inflection time (“Migration onset,” black square; see Methods section 5.2) for each of the three starting colony conditions. **E.** Estimated time of migration as determined by the inflection points from individual curves (as shown in D) for all data in C. Comparisons of migration timing between each condition were tested for significant differences using the Kruskal Wallis test (p-value=5.52×10^-29^) followed by Mann-Whitney post-hoc test (with Holm-Bonferroni method). The post-hoc test revealed that there was no significant difference between the two 2D conditions (p-value =0.13), however, each of the 2D conditions was different from the 3D lumenoid condition (*2D colony EMT* vs. *3D lumenoid EMT*: p-value =9.25×10^-26^; *2D PLF colony EMT* vs. *3D lumenoid EMT*: p-value =6.69×10^-18^). Significant differences with p-values <.001 were designated with *** and p-value ≥.05 were considered not significant (n.s.). **F.** 3D rendered example showing nuclear centroids inside (yellow spheres) and outside (cyan spheres) the basement membrane (BM, semi-transparent grey shell). Yellow and cyan spheres show nuclear centroids, not segmentations. **G.** Fraction of nuclei with centers outside of the basement membrane for each *3D lumenoid EMT* time-lapse where a nuclear marker was expressed at the time of migration (this included all *3D lumenoid EMT* time-lapses of mEGFP-tagged H2B and mEGFP-tagged Eomes, n=35) over time. The line is the median and the shaded region is the 25^th^-75^th^ percentile range **H.** An individual trajectory from G showing time of migration (black square) as determined by inflection point for the fraction of nuclei centered outside of the basement membrane (n=35 Methods section 5.3). **I.** Migration timing based on the fraction of nuclei outside the basement membrane in G compared to the method based on area at the glass in C (n=35).

We found that in *2D colony EMT* and *2D PLF colony EMT*, the colony area typically decreased prior to migration, corresponding to the initial contraction phase and nascent lumenoid formation observed previously (Fig. 3B and C), while in *3D lumenoid EMT*, the “footprint area” only increased slightly prior to cell migration (Fig. 5C and D). Next, across all conditions, a rapid rise in area occurred as cells began to migrate and fill the imaged field of view (Fig. 5C). For each time-lapse, we defined the time of migration as the inflection point via an automated computational approach (Methods section 5.2) that identified the start of this rapid rise in area (see black squares in Fig. 5D). We found that migration occurred earlier in the two 2D EMT conditions than in *3D lumenoid EMT*. The median migration timing for *2D colony EMT* (22.5 h [min-max:17.5-28.5 h]) and *2D PLF colony EMT* (23 h [min-max: 18.5-30 h]) conditions were significantly earlier than the median migration timing for *3D lumenoid EMT* (29.5 h [min-max: 21-38 h]; see Fig. 5E and Methods section 5.2 for statistical details). The migration timing of the two 2D conditions and the 3D lumenoid condition had a statistically significant difference (Kruskal-Wallis, p-value =5.52×10^-29^, see Fig. 5E for post-hoc test details and Methods section 5.2 for statistical method details) (Fig. 5E). The difference in migration timing from 2D and 3D conditions was still significantly different when each cell line used in this study was considered individually despite small differences in timings between the lines (Supplemental Fig. S3).

While migrating cells from 2D colonies primarily migrated across the glass, cells from 3D lumenoids could migrate both along the outside of the 3D lumenoid as well as across the glass (see Fig. 4A). This raised the possibility that the difference in migration timing that we identified using the footprint area of the lumenoid at the glass was due to a difference in where cells initiated migration, at the glass surface (in both 2D EMT conditions) vs. along the external wall of the basement membrane above the glass (only possible in *3D lumenoid EMT*). To address this, for each 3D lumenoid that expressed nuclear markers at the time of migration (mEGFP-tagged H2B and mEGFP-tagged Eomes, 35 movies total) we generated 3D instance segmentations of nuclei and a mesh representation of the basement membrane (Methods section 4.5, Supplemental Fig. S4, Fig. 5F). We measured the fraction of nuclei detected outside the basement membrane over time (Fig. 5G). We found that the fraction of nuclei outside was nearly zero until it rose rapidly with many migrating cells exiting the basement membrane (Fig. 5G). We applied the same inflection point analysis described above to detect migration timing (Fig. 5H). We found that migration timing calculated using this 3D approach (Fig. 5I and Methods section 5.3) (median 29 h [min-max: 21.5-33 h]) was very similar to the migration timing detected using the area at the glass metric for the same 35 *3D lumenoid EMT* movies (median 31 h [min-max: 24.5-35 h]). Although the 3D method consistently detected migration ∼2 h earlier than the area at the glass method, this delay was much less than the delay between 2D and 3D conditions, supporting the difference in migration timing found between 2D and 3D conditions.

### The expression timings of canonical EMT markers parallel the differences in migration timing found between 2D colony and 3D lumenoid EMT conditions

Since the timing of cell migration was later in *3D lumenoid EMT* compared to both 2D EMT conditions, we wondered whether the timing of expression of other markers would also be delayed in *3D lumenoid EMT* relative to the 2D EMT conditions. We investigated SOX-2, a protein marker of stemness that turns off with cell differentiation^24,25^ as well as Eomes and E-cadherin, protein markers known to change with EMT. Eomes is a T-Box transcription factor associated with mesodermal cell fate that turns on and then off with EMT^9,26^ and E-cadherin is known to be highly expressed in epithelial cells and downregulated with EMT^11,27–30^. To quantify how protein expression changed over time, we generated cell lines expressing mEGFP-tagged versions of these markers, imaged these cells during the three EMT conditions, and then measured fluorescence intensities within the all-cells masks over time.

Initial SOX-2 fluorescence intensities were different between *2D PLF colony EMT, 2D colony EMT, and 3D lumenoid EMT* (Fig. 6A, B, and C). However, it is difficult to determine whether these differences reflect a biologically meaningful effect of cell culture geometry on SOX-2 expression versus some other cause since we typically see that thicker samples appear brighter than thinner ones in most fluorescent cell lines that we image (data not shown). Instead, we compared how SOX-2 was lost across all conditions (Fig. 6B) by measuring the time at half maximal loss of SOX-2 for each trajectory (Fig. 6C). We observed that the timing of SOX-2 expression loss was similar across all conditions (Kruskal-Wallis, p-value = .45; see T_half-max_ in Fig. 6D). However, there was a delay in the migration time of *3D lumenoid EMT* compared to the 2D EMT conditions in the SOX-2 line (see T_migration_ in Fig. 6D) consistent with that shown across all the data (Fig. 5E). The similar timing in SOX-2 expression loss compared to the differential migration timing suggests that the observed loss of stemness is not related to EMT migration timing.

**Fig. 6:**
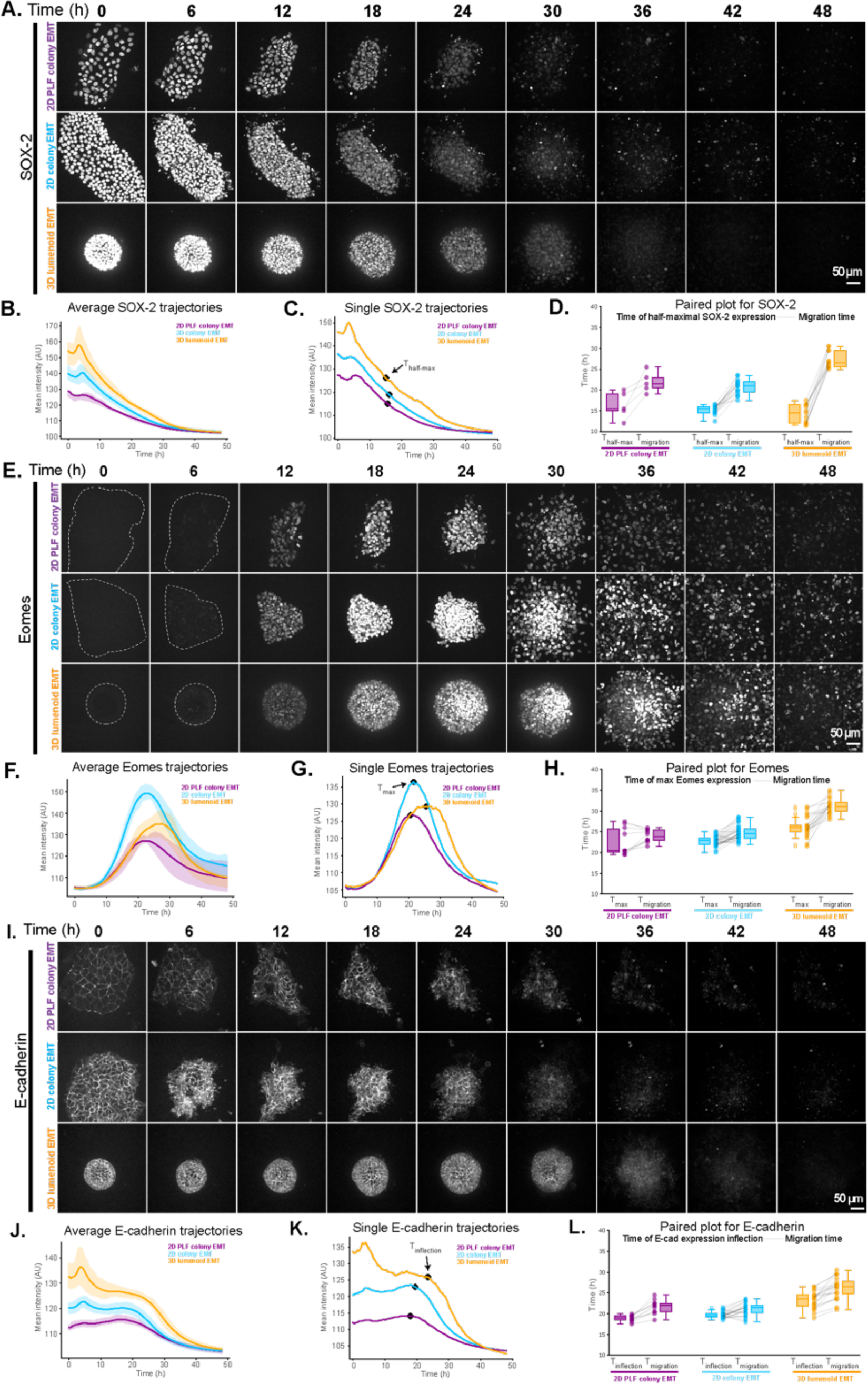
Fluorescence in bright-field-generated all-cells masks of EMT markers, but not SOX-2 show similar qualitative dynamics to that of migration timing. **A.** XY maximum intensity projections of mEGFP-tagged SOX-2 in a representative example from *2D PLF colony EMT* (top row), *2D colony EMT* (middle row), and *3D lumenoid EMT* (bottom row). Scale bar: 50 µm. **B.** Averages of all SOX-2 trajectories over time grouped by condition. Shaded regions show the 25^th^-75^th^ percentile distribution. **C.** Decrease in average SOX-2 fluorescence intensity over time for single trajectory examples with the time of half-max fluorescence intensity indicated (arrow). **D.** Time when SOX-2 fluorescence in all-cells masks reaches half the max value paired with time of migration for each condition. Matched pairs of time points from the same sample are connected with gray lines. *3D Lumenoid EMT* conditions have significantly later migration times than either *2D PLF colony EMT* (p=.01) or *2D colony EMT* (p=1.20×10^-5^) conditions, but not between the two 2D conditions (p=.41) according to Mann-Whitney post-hoc tests (Holm-Bonferroni p-value correction). Differences in timing of half-max fluorescence intensities were not statistically significant (Kruskal-Wallis, p-value=.45). Sample sizes are 5, 17, and 13 for *2D PLF colony EMT*, *2D colony EMT*, and *3D lumenoid EMT*, respectively. **E.** XY maximum intensity projections of mEGFP-tagged Eomes in a representative example from *2D PLF colony EMT* (top row), *2D colony EMT* (middle row), and *3D lumenoid EMT* (bottom row). The dashed white line outlines the perimeter of the colony or lumenoid drawn manually from the brightfield image. Scale bar: 50 µm. **F.** Averages of all Eomes trajectories over time grouped by condition. Shaded regions show the 25th-75th percentile distribution. **G.** Average Eomes fluorescence intensity over time for single trajectory examples with the time maximum fluorescence intensity indicated (arrow). **H.** Time of max fluorescence paired to time of migration for each time-lapse for each condition. Migration timing is significantly later in *3D lumenoid EMT* compared to either the *2D PLF colony EMT* (post-hoc p=1.9×10^-5^) or the *2D colony EMT* (p=4.38×10^-9^) conditions but not between 2D conditions (p=.21) according to Mann-Whitney post-hoc tests (Holm-Bonferroni p-value correction). Similarly, the time of max Eomes fluorescence is significantly different between *3D lumenoid EMT* and *2D PLF colony EMT* (p=.03), and between *3D lumenoid EMT* and 2*D colony* EMT (p=3.00×10^-6^) conditions but not between 2D conditions (p=.39) according to Mann-Whitney post-hoc tests (Holm-Bonferroni p-value correction). Sample sizes are 10, 32, and 21 for *2D PLF colony EMT*, *2D colony EMT*, and *3D lumenoid EMT*, respectively. **I.** XY maximum intensity projections of mEGFP-tagged E-cadherin in a representative example from *2D PLF colony EMT* (top row), *2D colony EMT* (middle row), and *3D lumenoid EMT* (bottom row). Scale bar: 50 µm. **J.** Averages of all E-cadherin trajectories over time grouped by condition. Shaded regions show the 25th-75th percentile distribution. **K.** Average E-cadherin fluorescence intensity over time for single trajectory examples with the inflection time indicated (arrow). **L.** Time of E-cadherin fluorescence inflection paired with time of migration for each time-lapse for each condition. All three conditions were significantly different from one another in time of E-cadherin fluorescence inflection according to Mann-Whitney post-hoc tests (Holm-Bonferroni p-value correction) (*2D PLF colony EMT* vs*. 2D colony EMT* p=.01, *2D PLF colony EMT* vs. *3D lumenoid EMT* p=3.40×10^-4^, *2D colony EMT* vs. *3D lumenoid EMT* p=3.00×10^-6^) and between 2D and *3D lumenoid EMT* conditions for time of migration (*2D PLF colony EMT* vs. *3D lumenoid EMT* p=2.82×10^-4^, *2D colony EMT* vs. *3D lumenoid EMT* p=2.00×10^-6^) but not between 2D conditions (*2D PLF colony EMT* vs. *2D colony EMT* p=.36). Sample sizes are 11, 23, and 20 for *2D PLF colony EMT, 2D colony EMT* and *3D lumenoid EMT,* respectively. See Supplemental Table 1 for web-based 3D Volume Viewer links for each time-lapse shown.

Eomes, a protein marker more typically associated with EMT, is known to rise and fall over the course of EMT^9,26^. As expected, Eomes rose and fell in intensity across all EMT conditions (Fig. 6E, F and G). We observed large differences in the magnitude of peak Eomes intensity between EMT conditions (Fig. 6F and G) but due to the difficulty of interpreting intensity amplitudes (as discussed above), we focused on the timing of the maximum peak in the intensity profile as the metric for this marker and a comparison point (Fig. 6G.). We found that the time of maximum Eomes intensity from cells in *3D lumenoid EMT* condition occurred significantly later than either of the 2D EMT conditions (Kruskal-Wallis, p-value = 1.18×10^-5^, see Fig. 6H for post-hoc test details and Methods section 5.4 for statistical details), although there was also a large amount of variation in this timing for *2D PLF colony EMT* (Fig. 6F and T_max_ in H). Interestingly, both the timing of peak Eomes and that of migration were delayed in *3D lumenoid EMT* compared to the 2D EMT conditions (Kruskal-Wallis, p-value = 9.41×10^-10^, see T_migration_ and post-hoc test details in Fig. 6H and Methods section 5.4 for statistical details).

We also investigated the dynamics of E-cadherin, a known protein marker associated with EMT that is not a transcription factor. E-cadherin, which is highly expressed in epithelial cells, is downregulated with EMT^11,27–30^. As expected, we found that E-cadherin fluorescence intensity was lost across all conditions, rapidly decreasing between 20 and 30 h after EMT induction (Fig. 6I, J and K). To compare the timing of E-cadherin loss between conditions, we identified the inflection point just before the rapid decrease in E-cadherin intensity (Fig. 6K). We saw that there were statistical differences in the timing of E-cadherin loss between all EMT conditions (Kruskal-Wallis, p-value = 5.5×10^-8^), however, the differences between 2D and 3D EMT conditions were much larger than that between the two 2D conditions (see T_inflection_ and post-hoc test results in Fig. 6L). As before, there remained a significant delay in migration timing in 3D lumenoid EMT relative to the 2D EMT conditions (Fig. 5E and T_migration_ in Fig. 6K). Therefore, similar to Eomes, both the loss of E-cadherin and the timing of migration were delayed in 3D lumenoid EMT compared to 2D EMT conditions. However, in this case, the 2D conditions had slightly significantly different timings of E-cadherin loss and similar migration timings (see T_migration_ and post-hoc test results in Fig. 6L).

The interpretation of the relative timing of these expression changes more generally across the entire dataset is made difficult by the observed variability in migration timing between cell lines (Supplemental Fig. S3). For example, the loss of E-cadherin expression (via the inflection point) occurs earlier than the peak of Eomes expression (Supplemental Fig. S5A), but cell migration is also earlier in the E-cadherin line compared to the Eomes line (Supplemental Fig. S3). If instead we plot these expression dynamics relative to the migration time, either by absolute comparison (Supplemental Fig. S5B) or normalized between the time of induction and migration (Supplemental Fig. S5C), there is no longer a discrepancy in the timing of the Eomes expression peak and E-cadherin expression loss, implying that these occur at similar times relative to eventual migration.

### mEGFP-tagged ZO-1 reveals organizational changes in hiPS cells undergoing EMT, including the contraction of apical faces and cell delamination

The loss of E-cadherin expression also suggested a change in the make-up or presence of the cell-cell junctions where E-cadherin localizes and functions. Changes in composition and later loss of cell-cell junctions are hallmark organizational changes that occur over the course of EMT. Cadherins, components of adherens junctions, are known to regulate the adhesiveness between cells^31,32^ and to change in composition over the course of EMT^33^. In addition to observing changes in cell junction composition with E-cadherin, we also investigated the loss of cell-cell junctions using ZO-1. ZO-1 is a component of tight junctions, which are responsible for the barrier-forming function of epithelial cells^34^. Tight junctions are lost with EMT as cells lose their apicobasal polarity^35,36^.

The tight-junction network and its loss over the course of EMT can be seen in the time-lapse imaging of an mEGFP-tagged ZO-1 hiPS cell line (Fig. 7A). To quantify the changes in fluorescence intensity and localization of ZO-1 we generated all-cells masks from the bright-field channel of these movies and measured the average ZO-1 fluorescence intensity over time as a function of the height above the glass (Fig. 7B). In *2D PLF colony EMT*, the ZO-1 signal remains at a similar height above the glass up until it is lost after the onset of migration (Fig. 7B left). In contrast, in *2D colony EMT,* the ZO-1 signal increases in height dramatically in the period prior to migration from ∼14 h - 22 h (Fig. 7B middle). After migration onset, some ZO-1 fluorescence re-localizes to a lower Z-plane (black arrow in Fig. 7B middle), and the overall fluorescence intensity in both Z-locations then decreases in the following hours. Interestingly, in *3D colony EMT* we observe a similar initial increase in ZO-1 height that is then followed by a striking accumulation of fluorescence intensity at low Z-planes (black arrow in Fig. 7B right), which persists for ∼4 h before decreasing after migration onset (Fig. 7B right). These patterns of ZO-1 expression and localization were very reproducible (Fig. S6).

**Fig. 7:**
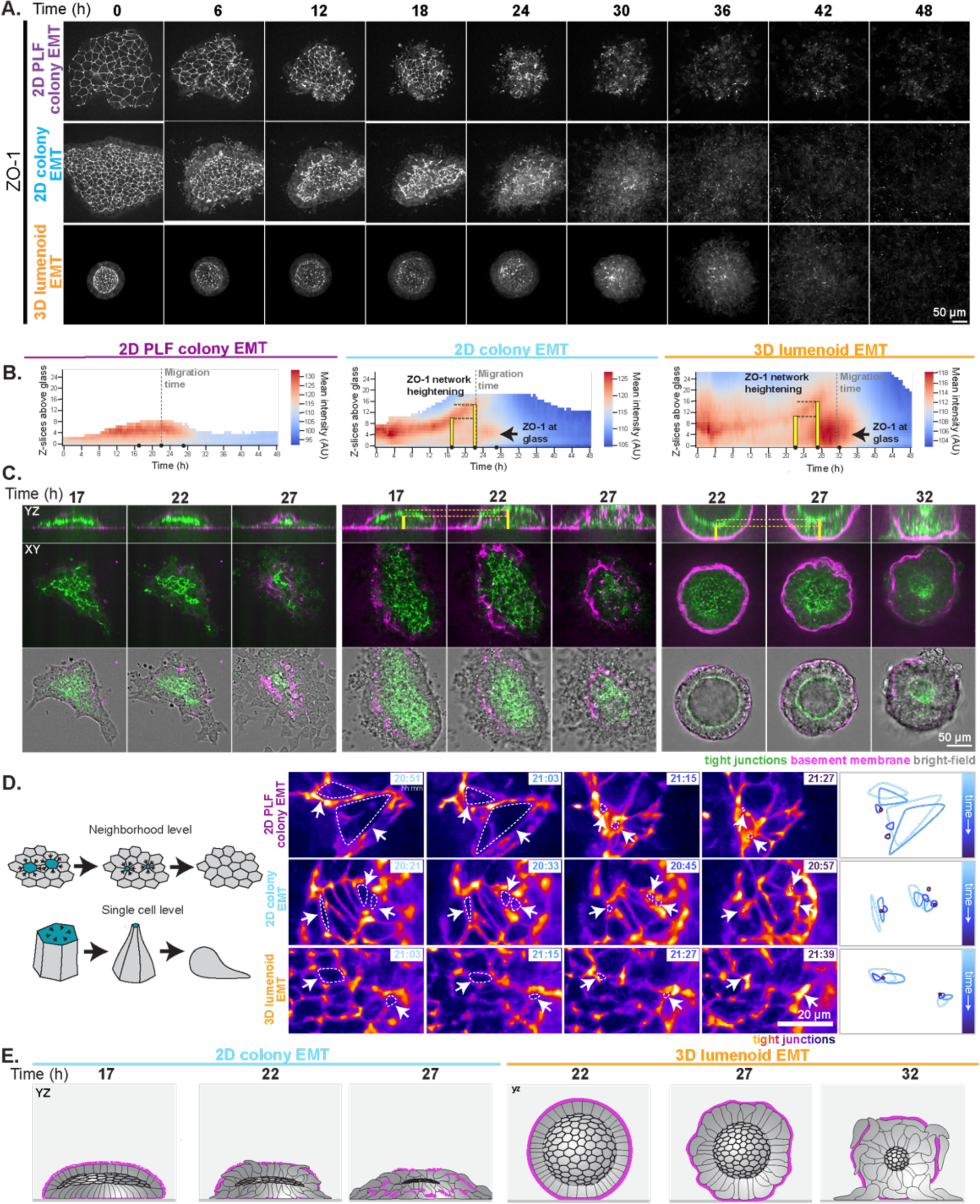
hiPS cells undergoing EMT constrict their apical faces to a point before delaminating in all three hiPS cell-EMT conditions. **A.** Maximum intensity Z-projections of mEGFP-tagged ZO-1 in 2D EMT conditions (top two rows) and *3D lumenoid EMT* (bottom row). Scale bar: 50 µm. **B.** Heatmaps showing the mean ZO-1 fluorescence intensity within the all-cells masks as a function of Z-height above the glass and over time for example time-lapses from each EMT condition. The migration times for the given examples are indicated by the vertical, gray dashed lines. The increase in Z of the bright ZO-1 fluorescence (red) over time shows the ZO-1 network heightening – compare yellow bars indicated in B and C. Black arrows in *2D colony EMT* and *3D lumenoid EMT* plots show where ZO-1 fluorescence accumulates at the glass around the time of cell migration. Black dots on the time axis indicate the times at which images from the same time-lapse were selected and shown in C. **C.** XY maximum intensity projections of five Z-slices and YZ maximum intensity projections through approximately the same thickness in Y (50 pixels) centered about the ZO-1 network. Yellow bars drawn from the glass to the top of the network are drawn in the YZ view of the 17 and 22 h time points in *2D colony EMT* and 22 and 27 h time points in *3D lumenoid EMT* to highlight the heightening of the ZO-1 network prior to migration. The three time points shown for 3D lumenoid EMT are shifted later than those shown in the 2D EMT examples because migration occurs later from 3D lumenoids than 2D colonies. **D.** Left: Schematic of neighborhood level changes in shapes of apical faces and singe cell level changes in cell shape expected with delamination. Right: Example cropped regions of the ZO-1 network colored by ZO-1 intensity that show constriction of apical faces to a point within each EMT condition. Images are maximum intensity projections of two slices of denoised data shown in pseudo-color (see Methods section 4.3). Dotted lines follow the shape of the apical face of the same cell in each time point. Overlay of the dotted lines color-coded by time point are shown on the right. Scale bar: 20 µm (see Supplemental Movies 4-6). **E.** Schematic of proposed model showing the effect of the different basement membrane environments in *2D colony EMT* vs. *3D lumenoid EMT* on the relationship between the timing of delamination and cell migration. See Supplemental Table 1 for web-based 3D Volume Viewer links for each time-lapse shown.

Three time points are shown in Fig. 7C that illustrate the key changes in ZO-1 patterns observed in Fig. 7B (compare YZ views in the top row of Fig. 7C with corresponding time points in Fig. 7B). These time points show examples of the ZO-1 network in the context of where the cells are (given by bright-field) and the basement membrane (given by anti-collagen IV labeling) surrounding the time of cell migration. The ZO-1 network appears to shrink between the first two time points, given by a decrease in the area of the patch of ZO-1 in the 2D EMT conditions, and a decrease in the inner diameter of the lumen in *3D lumenoid EMT*. At the same time that the ZO-1 network shrinks, a thickening of the bottom layer of cells in *2D colony EMT*, and a thickening of the lumenoid wall in *3D lumenoid EMT* occurs (see the comparison of the heights indicated by the yellow bars in Fig. 7B and C), suggesting that cells are becoming taller and/or potentially losing their connection with the ZO-1 network and delaminating from the epithelial-like sheet. By the final time points, the ZO-1 network is greatly reduced and many cells seem to have delaminated. Interestingly, cells appear to begin migrating in the 2D conditions at the same time that the ZO-1 network shrinks (22 h panel in 2D conditions in Fig. 7C), while cells migrate later than ZO-1 network shrinkage in *3D lumenoid EMT* (27 h panel for network shrinkage and 32 h panel for cell migration in 3D lumenoid condition in Fig. 7C). Together, these results suggest a decoupling in the timing of cell delamination from the ZO-1 sheet and subsequent cell migration time in *3D lumenoid EMT*, but not in the 2D EMT conditions.

To examine delamination more closely, we looked at changes in the apical faces of cells over time. Shapes of apical faces and single cells often change during EMT due to apical constriction and cell delamination (Fig. 7D left). For example, during gastrulation, ingressing cells have been shown to proceed through this type of mechanism^37–41^. Across all three of the EMT conditions, we observed that the apical faces of cells, as outlined by ZO-1, became smaller on average up until cell migration onset (Fig. 7C). Upon closer inspection using higher temporal imaging (every 3 min), we observed that the apical faces typically shrink to a point before disappearing entirely (Fig. 7D right and Supplemental Movies 4-6), suggesting that cells in all three EMT conditions do indeed leave the epithelial sheet via apical constriction and delamination. Apical face closure could be detected by 22 h post EMT induction in all conditions, and often multiple cells in a local neighborhood would close their apical faces simultaneously (Fig. 7D).

Together, the constriction of apical faces to a point, the shrinkage in the ZO-1 network size, the thickening of lumenoid walls, and the delayed migration onset relative to these events in *3D lumenoid EMT* led us to propose the following model: In 2D EMT conditions, cells are able to begin migrating as soon as delamination occurs because the basement membrane is not well-established. In contrast, cells in the *3D lumenoid EMT* condition do not migrate until after delamination occurs because they are trapped inside the more mature basement membrane (Fig. 7E).

### Different plating conditions and cell geometries display different degrees and patterns of basement membrane lift-off from the glass during EMT

In addition to changes at the apical faces of cells, we reasoned that associations with the basement membrane on the basal side of cells could also influence 2D colony or 3D lumenoid shape. We found that the basement membrane at the glass was relatively unchanged in *2D PLF colony EMT* over the course of imaging, possibly due to this more adhesive substrate effectively holding it in place (Fig. 8A). However, in *2D colony EMT*, the basement membrane appeared to become fragmented and was lifted off the glass (Fig. 8B). This coincided with the onset of cell delamination and migration.

**Fig. 8:**
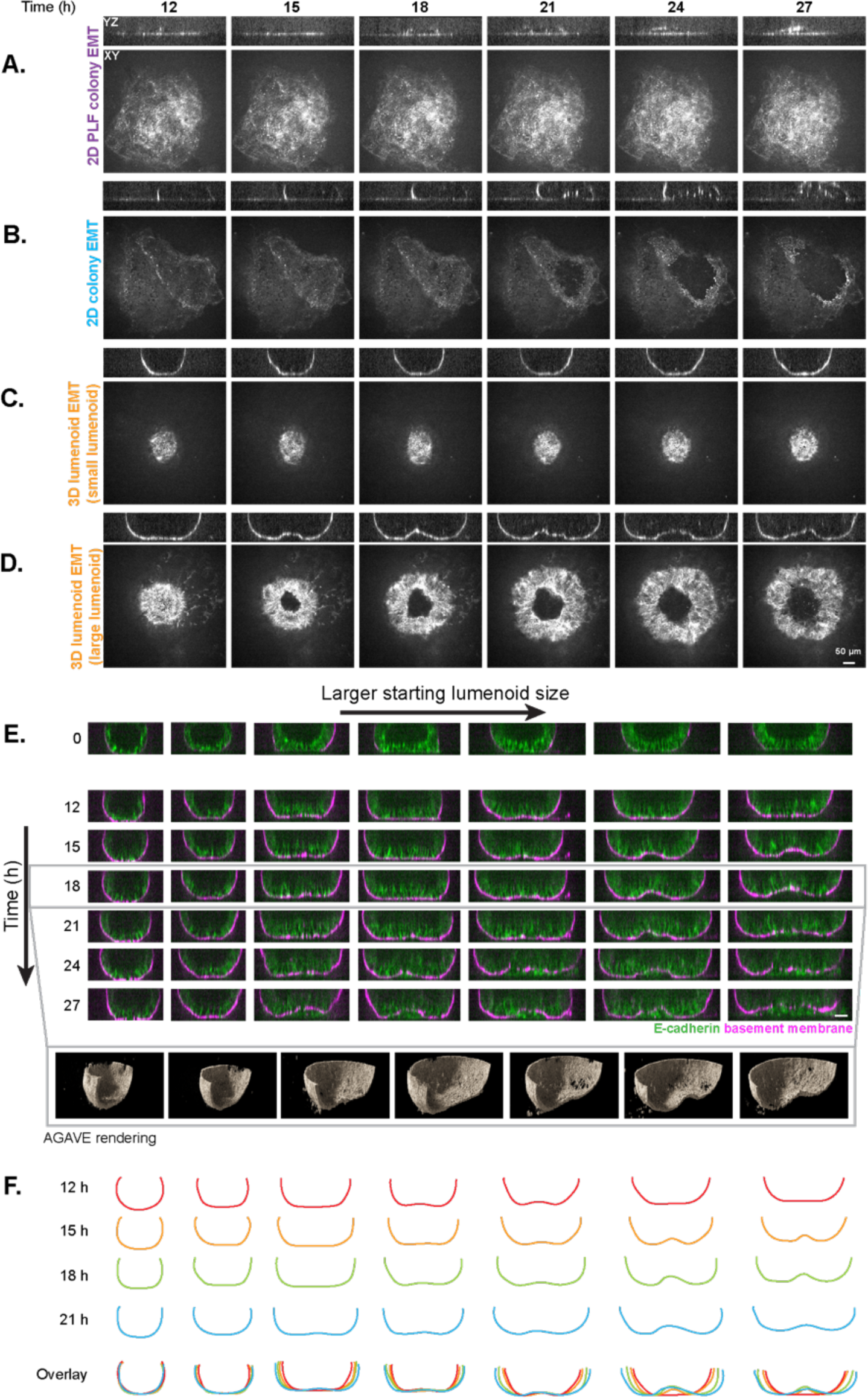
Basement membrane detachment from the glass depends on the adhesiveness of the substrate in 2D colonies and size of the lumenoid in 3D. **A-D.** Single slices in XY at the glass surface and in YZ cross section showing fluorescent collagen IV antibody in *2D PLF colony EMT*, *2D colony EMT*, and *3D lumenoid EMT* conditions over time. Scale bar: 50 µm. **A.** The basement membrane is not lifted from the glass in *2D PLF colony EMT*. **B.** The *2D colony EMT* condition lifts patches of basement membrane over time. **C.** Small lumenoids in the *3D lumenoid EMT* condition maintain their footprint on the glass prior to cell migration. **D.** Large lumenoids in the *3D lumenoid EMT* condition typically lift a single coherent dome of basement membrane, or a “dimple”, in the center of their footprint on the glass prior to cell migration. **E.** XZ side-views of lumenoids over time from the *3D lumenoid EMT* condition from smallest (left) to largest (right) expressing mEGFP-tagged E-cadherin (green) and labeled with a fluorescent collagen IV antibody (“basement membrane,” magenta). Scale bar: 50 µm. AGAVE renderings of the basement membrane from the lumenoids at the 18 h time point, cut in half to show dimple formation at the center. **F.** Manual tracing of the basement membrane in the side-views in E reveals a similar curvature for small lumenoids without dimples and larger lumenoids with dimples. See Supplemental Table 1 for web-based 3D Volume Viewer links for each time-lapse shown.

In *3D lumenoid EMT*, we observed a striking size-dependent phenomenon where small lumenoids displayed little change in their basement membrane footprint on the glass (Fig. 8C), whereas large lumenoids lifted off the glass in the center of their footprint, forming a coherent dome, or “dimple,” often well before the onset of migration (Fig. 8D). Without EMT induction, we did not observe dimple formation in small or large lumenoids (Fig. 8E, T=0 h). However, with EMT induction, dimple formation was very robust and highly size dependent: Dimples formed earliest in large lumenoids, later in mid-sized lumenoids, and not at all in small lumenoids (Fig. 8E). The curvature of small lumenoids was similar to that generated by the dimples formed in larger lumenoids, suggesting that there may be a preferred curvature for lumenoids preparing to undergo EMT, or perhaps a buckling wavelength of the pre-EMT epithelium (Fig. 8F).

## Discussion

EMT is a fundamental cell state transition in contexts ranging from healthy embryonic development to disease states. The study of this dynamic process is challenging due its diverse characteristics in different biological contexts and widely varying definitions based on cell function, organization, molecular census and/or environment. We introduce an hiPS cell model system for developing a holistic understanding of EMT across all these fundamental cell properties. We previously demonstrated the power of hiPS cells grown in 2D epithelial-like cultures as a model for characterizing cell organization across multiple labeled structures, demonstrating the potential for data integration across one type of cell observation in a naturally immortal and karyotypically normal human cell type^42^. Here, we demonstrate how taking advantage of an hiPS cell model system in the dynamic context of differentiation offers a robust system for gaining a holistic understanding of EMT across multiple types of observations in cells and geometrical contexts.

The hiPS cell-EMT process is robust to cell culture geometry, reproducibly inducible in both 2D and 3D conditions. 2D cell cultures are typically more experimentally tractable and amenable to live imaging, but cells are often forced into an artificially flat arrangement and cells at of the center of the colony can exhibit different organization and gene expression than those at the edge^42^. On the other hand, while cells studied *in vivo* are in their native 3D arrangement, these assays are often challenging to perform and interpret due to the complexity of the cell composition and environment (i.e. diversity of cell types and chemical signaling gradients) and inaccessibility of the system. There is therefore a need for a cell culture system that moves towards the physiological relevance of a 3D cell culture geometry while optimizing the experimental manipulability of an *in vitro* assay^43^; we present the adherent 3D hiPS cell lumenoid as such a system. All cells in the lumenoid have uniform polarity (with apical sides inwards and basal sides out), with a central lumen providing a shared signaling environment. The environment of 3D lumenoids is easily accessible and controllable since only a dilute concentration of Matrigel is needed to form the 3D structure, unlike most 3D cell culture models that require embedding in a gel for mechanical support. This enabled us to easily include a human specific, anti-collagen IV antibody to visualize the basement membrane in live human cells. Furthermore, we leveraged the flexibility of the hiPS cell model system to be cultured as either 2D colonies or 3D lumenoids to perform direct comparisons between multiple geometrical contexts in the same model system.

The high reproducibility of EMT in this model system allowed us to leverage a wide array of fluorescently tagged cell lines all undergoing the same transition so that we could integrate multiple types of observations in cells. First, we looked at the relationship between changes in marker expression and migration timing. The timing of expression changes in EMT-related markers Eomes (based on the time of peak fluorescence intensity) and E-cadherin (based on the inflection before rapid loss of fluorescence) were delayed in 3D lumenoid EMT compared to 2D EMT conditions. The timing of migration onset was similarly affected by initial hiPS cell culture geometry. In contrast, changes in stemness marker SOX-2 (based on the time of half-maximal decrease in fluorescence) was not different between the cell culture geometries, but there was still a delay in migration onset in 3D lumenoid EMT compared to the 2D EMT conditions. Together, these results suggest there could be a relationship between EMT-related markers and migration timing while the loss of SOX-2 is unrelated to migration timing. It is important to recognize that this study does not elucidate the exact order or causality between the different changes we observed over the course of EMT. Additionally, the causality may include other unexplored factors such as the role of cell death and mitosis on the timing of cell migration (and the course of EMT in general). These processes are known to be critical for maintaining the number of cells in an epithelial sheet and are, in the case of mitosis, important for the timing of cell ingression^44^. We hope that the large, annotated, standardized, and publicly available dataset produced from this study will provide a rich resource for future discovery to addressing questions like these.

We hypothesize that the difference in migration timing between 2D and 3D EMT conditions may be due to differences in the basement membrane environment of the different EMT conditions: A new 3D basement membrane shell does form in 2D EMT conditions if Matrigel is included during EMT induction, as the colony forms a nascent lumenoid prior to cell migration. However, this shell appears weaker (dimmer fluorescence intensity) than that of the preformed 3D lumenoid in the *3D lumenoid EMT* condition. Therefore, the barrier to cross the basement membrane may be lower in 2D EMT conditions than in *3D lumenoid EMT*. In addition to the differences in the barrier-forming strength of the basement membrane to resist cell migration in 2D vs. 3D cell culture geometries, labeling the basement membrane provides other insights. We saw evidence, at least in the 3D context, of mechanical displacement of the basement membrane during cell migration via the brightening of collagen IV labeling around the edges of holes formed by migrating cells (analogous to how pulling apart window curtains collects fabric at the window edge), similar to what has been seen in vivo^40,45^. However, how mechanical displacement balances among the multiple known mechanisms by which the basement membrane is altered by cells during EMT remains to be determined, including the relative contributions of protease digestion by MMPs, the mechanical opening of holes by cells migrating through the sheet, and mechanical pulling on the sheet by migrating cells^46^. Future studies to manipulate the integrity of the basement membranes of the 2D and 3D cell culture geometries in hiPS cells (e.g. the use of MMP inhibitors and collagenase) are needed to determine to what extent each mechanism of basement membrane remodeling is employed in the different geometrical contexts. In addition, future inspection, especially with high-resolution microscopy, of the de-novo deposition of basement membrane during 3D lumenoid formation from 2D colonies may also reveal insights into the process and mechanism by which cells change between these cell culture geometries.

We also provide an example of the use of the 3D lumenoid hiPS cell culture geometry on its own to study an example of mechanical properties that change with EMT. The results that show dimpling in the epithelial sheet of the 3D lumenoid prior to cell migration in the *3D lumenoid EMT* condition exemplify the advantage of a uniform mechanical environment to study mechanical properties that change during EMT that would be much harder (if at all possible) to study and interpret in a 2D system. The ability of lumenoids to form dimples indicates that the lumenoid basement membrane is both strong and flexible, maintaining its integrity even as it is lifted off the glass, and that by spanning many cell lengths this structure enables individual cells to act as a coherent, mechanical collective. The size threshold of dimple formation suggests that apical closure could be driving a preferred curvature of the lumenoid. However, there are many other mechanisms that likely also contribute to this effect and are not mutually exclusive. Epithelial buckling in particular has been linked to cell growth, area confinement, cell shape changes, and changes in cell-cell adhesion^47^ – all of which are occurring in the lumenoid EMT system. Furthermore, modeling of 2D epithelia suggests that a contractile ring spanning many cell lengths can cause a 2D epithelium to bend out of plane in a configuration that looks remarkably similar to the lumenoid dimple ^48^, and we do in fact observe apical cell-cell junctions encircling the dimple in large lumenoids using the ZO-1-tagged line (data not shown). It is also possible that an entirely different type of mechanism could be at play, such as accumulation of solutes below the lumenoid driving dimple formation.

These different types of observations highlight the value of comparing and integrating insights gained from studying EMT across cell structures and culture geometries to gain a holistic view of this essential and ubiquitous, yet thus-far poorly defined, biological process. With its ability to provide these insights with incredible reproducibility, the hiPS cell model system offers a tractable and robust system with which to quantitatively probe the dynamic changes in cell function, organization, molecular census, and environment that are hallmarks of the EMT process. This approach, together with the wealth of time-lapse imaging data and easily accessible analysis tools used to produce these results, presents a valuable resource for future work in this field.

## Acknowledgements

We thank Victoria Hurless, Clare Gamlin and Irina Mueller for program management support, Sean Meharry and Daniel Toloudis for software infrastructure support, Chris Frick and Erin Angelini for technical and literature support, Hannah Thorp for dataset review and the Allen Institute for Cell Science team for helpful discussions and support. We thank Javier Chávez, Lauren M. Kuo, Jacob McCarley, Jacqueline E. Smith, Danielle Yi for cell line production. Ben Z. Stanger, M. Angela Nieto, Magdalena Zernicka-Goetz, and Christine L. Mummery for helpful scientific discussions. The figures from this manuscript have been prepared using the checklist from Schmied et al^49^. The image data and metadata shared in this study have been prepared by following the REMBI guideline^50^. The WTC-11 hiPS cell line used to create the gene-edited cell lines of the Allen Cell Collection was provided by the Bruce R. Conklin Laboratory at the Gladstone Institute and UCS. J.A.T. was supported by the Howard Hughes Medical Institute. This article is subject to HHMI’s Open Access to Publications policy. HHMI lab heads have previously granted a nonexclusive CC BY 4.0 license to the public and a sublicensable license to HHMI in their research articles. Pursuant to those licenses, the author-accepted manuscript of this article can be made freely available under a CC BY 4.0 license immediately upon publication. We wish to thank Allen Institute founders, Jody Allen and Paul G. Allen, for their vision, encouragement and support.

## Author contributions according to Contributor Role Taxonomy (CRediT)

Conceptualization, C.H., A.B., L.K.H., G.D., N.N., J.A.T., S.M.R., R.N.G.; Writing – original draft, C.H. A.B., L.K.H., G.D., S.M., N.N., J.C.D., C.L.L., G.N., S.E.G., M.F.S., D.J.T.; Writing – review & editing, C.H., A.B., L.K.H., S.C., G.D., S.M., N.N., T.B., J.C.D., J.H.E., C.L.L., S.E.P., J.A.T., S.M.R., R.N.G.; Methodology, C.H., A.B., L.K.H., S.C., G.D., S.M., N.N., A.J.F., G.N., S.A.O., S.E.G., M.F.S., D.J.T., M.P.V.; Data curation, A.B., S.C., S.M., E.M.A., T.B., J.H.E., M.J.H., H.S.M., S.E.P., A.P., E.E.S., L.S.S., D.J.T., J.P.T.; Formal analysis, L.K.H., N.N., S.E.P., S.M.R.; Validation, C.H., A.B., L.K.H., S.C., S.M., N.N., G.N., S.A.O., M.F.S., D.J.T., M.P.V.; Visuzalition, C.H., A.B., L.K.H., S.C., N.N., J.H.E., S.E.P., M.F.S.; Investigation, C.H., A.B., L.K.H., S.C., S.M., E.M.A., T.B., J.H.E., A.J.F., M.J.H., H.S.M., G.N., S.A.O., S.E.P., A.P., E.E.S., M.F.S., L.S.S., D.J.T., M.P.V., J.A.T., S.M.R., R.N.G.; Project administration, C.H., A.B., J.G., S.M.R.; Resources, E.A.E., M.A.F., B.W.G., K.N.K., B.R., L.S.S., J.R.T., G.T., C.S.W., J.Y.; Software, S.M., P.G., S.L.M., G.N., M.F.S., D.J.T., M.P.V., L.W.; Supervision, C.H., A.B., L.K.H., G.D., B.R., M.P.V., J.Y., S.M.R., R.N.G.

## 1. Experimental methods

### 1.1 Cell lines and plasmids

All work with human induced pluripotent stem (hiPS) cell lines was approved by internal oversight committees and performed in accordance with applicable National Institutes of Health, National Academy of Sciences, and International Stem Society for Stem Cell Research guidelines. We used the parental WTC-11 hiPS cell line derived from a healthy male donor^51^ as the basis for developing each CRISPR-Cas9 genome edited line, which features an endogenous fluorescent protein linked to a specific cellular structure protein^52,53^. Along with the unedited WTC-11 line, we used the following cell lines: AICS-0061 cl.36 HIST1H2BJ-mEGFP which express mEGFP-tagged histone H2B type 1-J (H2B), AICS-0036 cl.6 AAVS1 mEGFP which express mEGFP from AAVS1 (localized to the cytoplasm and nucleus), AICS-0122 cl.4 EOMES-mEGFP which express mEGFP-tagged Eomesodermin (Eomes), AICS-0114 cl.32 CDH1-mEGFP which express mEGFP-tagged E-cadherin, AICS-0074 cl.26 SOX2-mEGFP which express mEGFP-tagged transcription factor SOX-2 (SOX-2) and AICS-0023 cl.20 ZO-1-mEGFP which express mEGFP-tagged tight junction protein ZO-1 (ZO-1). The Eomes and E-cadherin cell lines were newly generated in this study for the Allen Cell Collection, using methods with AAV6 DNA donors^54^. In brief, cells underwent nucleofection using a Lonza P3 Primary Cell 4D Nucleofector X Kit S (Cat. # V4XP-3032) and a Lonza 4D system (Lonza 4D Core Unit Cat. # AAF-1003B with Nucleofector X Unit Cat. # AAF-1003X) with Cas9/sgRNAs ribonuclear proteins, then transduced with custom designed AAV6 vectors at 0.5×10^3^ MOI^55^. Following gene editing, cells from the modified populations were identified and, where needed, enriched with custom-designed droplet digital (ddPCR) assay surveillance. Subsequently, PCR screening was used to identify clones. Comprehensive details regarding these cell lines are accessible via the Allen Cell Collection (https://www.allencell.org/), and cells can be acquired through Coriell (https://www.coriell.org). The donor plasmids essential for creating these cell lines can be obtained through Addgene (https://www.addgene.org/The_Allen_Institute_for_Cell_Science/).

### 1.2 Cell culture

Cells were cultured in mTeSR^TM^1 (STEMCELL Technologies, Cat. # 85850) supplemented with 1% penicillin/streptomycin (P/S) (Gibco, Cat. # 15070063), on surfaces pre-coated with growth factor-reduced Matrigel (Corning, Cat. # 356231). Media was changed daily. When the hiPS cells reached a confluency of 70-85% (every 3-4 days), they were dissociated into single cells using Accutase (Gibco, Cat. # A11105-01) and then re-seeded in supplemented mTeSR1 medium with 1% P/S and a 10 μM concentration of the ROCK inhibitor Y-27632 (STEMCELL Technologies, Cat. # 72308). A comprehensive protocol is available at https://www.allencell.org/.

### 1.3 2D and 3D cell culture on glass-bottom imaging plates

The hiPS cells were enzymatically dissociated using Accutase (Gibco, Cat. # A11105-01) and seeded on 96-well CellVis glass imaging plates (CellVis, Cat. # P96-1.5H-N) at a density of 750-2,000 cells per cm^2^ in mTeSR™1 (STEMCELL Technologies, Cat. # 85850) supplemented with 10 µM Y-27632 (STEMCELL Technologies, Cat. # 72307). Prior to cell seeding in glass-bottom plates, the glass surfaces were coated either with PLF or Matrigel. The PLF coating consisted of 50 µg/mL Poly-D-Lysine (Sigma-Aldrich, A-003-E) and a combination of 18 µg/mL recombinant human Laminin 521 (Biolamina, Cat. # LN521) and 5 µg/mL Fibronectin (Gibco, Cat. # 33016-015). The Matrigel coating consisted of a 1:52 dilution of growth factor-reduced Matrigel (Corning, Cat. # 356231) in DMEM/F12 (Gibco, Cat. # 21041025). These coated plates were then cured in a tissue culture incubator set at 37 °C for 1-2 h. Once seeded, cells were maintained in mTeSR^TM^1 media, replaced daily for four days for 2D conditions. To initiate 3D lumenoid formation, hiPS cells were cultured on these plates for two days. On the third day, the culture medium was replaced with mTeSR™1 medium that included a 1:60 dilution of Matrigel. The fourth day additional mTeSR™1 medium was added to the wells. For all conditions (*2D colony EMT*, *2D PLF colony EMT*, and *3D lumenoid EMT, 2D no Matrigel EMT*) EMT was induced on day five post seeding. To visualize the basement membrane, Fluor® 660-conjugated anti-collagen IV antibody (Thermo Fisher, Cat. # 50-9871-82) was added at a dilution of 1:600 on day five post seeding.

### 1.4 EMT induction

In all EMT conditions except for *2D no Matrigel EMT*, EMT was induced by substituting the mTeSR™1 medium with RPMI-1640 (Gibco, Cat. # 11835-030), enriched with 5 µM CHIR99021 (Cayman Chemical, Cat. # 13122), B27 supplement devoid of insulin (Gibco, Cat. # A1895601) with a 1:60 Matrigel dilution. The *2D no Matrigel EMT* condition did not receive the 1:60 Matrigel dilution.

## 2. Image acquisition

### 2.1 Imaging position selection

A 10x bright-field well overview was used to select 2D and 3D colonies for imaging to ensure consistent colony sizes throughout the dataset. Following 10x overview acquisition, plates were removed from the microscope and taken for EMT induction as described above. Plates were then placed back on the microscope for imaging within 15 min of induction.

### 2.2 Acquisition of 60-hour, 3D time-lapse movies at 30 min intervals

Imaging was performed on three identical ZEISS Axio Observer Z1 spinning-disk confocal microscopes with 10x/0.45 NA Plan-Apochromat (for well overview and position selection) and 20x/0.8 NA Plan-Apochromat (Zeiss) (for dataset collection) and ZEN 3.2 software (blue edition; ZEISS). These microscopes were equipped with a 1.2x tube lens adapter for a final magnification of 12x or 24x, respectively, a CSU-X1 spinning-disk scan head (Yokogawa) and two Orca Fusion BT cameras (Hamamatsu). Microscopes were outfitted with a humidified environmental chamber to maintain cells at 37 °C with 5% CO_2_ during imaging. Live imaging for 60 h every 30 min was performed using the following laser powers (measured with 10x objectives): 488 nm at 0.77 mW, and 638 nm at 0.8 mW. An Acousto-Optic Tunable Filter (AOTF) was used to simultaneously modulate the intensity of all laser lines. The following Band Pass (BP) filter sets (Chroma) were used: 525/50 nm for detection of the mEGFP tag, and 706/95 nm for detection of the fluorescent collagen IV antibody. Images were acquired with an exposure time of 50 ms. Transmitted light (bright-field) images were acquired using a red LED light source with peak emission of 740 nm with narrow range and a far-red emission filter 706/95 for bright-field light collection. Optical control images and laser power were acquired using Argolight slides (Argo-CHECK and Argo-POWER slides respectively) at the start of each data acquisition to monitor microscope performance. Optical control images included dual camera acquisition of the Argolight slide field of rings pattern and black reference images. Laser power linearity for each wavelength, laser alignment by hotspot-centering, and estimation of spherical aberration effects via imaging of a fluorescent sphere of defined size were performed monthly.

This methodology was used to aquire 88 mEGFP-tagged H2B, 38 cytoplasmic mEGFP, 94 mEGFP-tagged Eomes, 51 mEGFP-tagged SOX-2, 79 mEGFP E-cadherin, and 99 mEGFP-tagged ZO-1 60-hour time-lapse movies. A total of 139 2D PLF colonies, 160 2D colonies, and 150 3D lumenoid conditions were collected across all cell lines.

### 2.3 Acquisition of 3D time-lapse movies at three-minute intervals

Five additional movies were acquired on a faster, three-minute interval, timescale. Time-lapses were acquired using an Eclipse Ti2 Nikon spinning-disk confocal microscope with a 10x/0.45 NA (for well overview and position selection) and 20x/0.8 NA Plan-Apochromat air objective or a 20x/0.95 NA Apochromat water-immersion objective (for dataset collection) and NIS-Elements AR 5.24.04 software (Nikon). The confocal was equipped with a CSW-1 spinning-disk scan head (Yokogawa) and two Orca Fusion BT cameras (Hamamatsu). The microscope was outfitted with a humidified environmental chamber to maintain cells at 37 °C with 5% CO_2_ during imaging. Live imaging every 3 min for 15 or 48 h was performed using the following laser powers (measured with 10x objectives): 488 nm at 0.89 mW, and 638 nm at 1.39 mW. Images were acquired with an exposure time of 50 ms. Transmitted light (bright-field) images were acquired using a red LED light source with peak emission of 740 nm with narrow range and a far-red emission filter 705/72 (Chroma) for bright-field light collection. Laser power linearity was assessed monthly for each wavelength.

This methodology was used to acquire two mEGFP-tagged H2B and three mEGFP-tagged ZO-1 three-minute interval time-lapse movies. A total of one *2D PLF colony EMT*, one *2D colony EMT*, and three *3D lumenoid EMT* conditions were collected across all cell lines.

### 2.4 Refractive index mismatch correction

The effects of spherical aberrations on Z-step size accuracy resulting from the refractive index mismatch between the air objective and the culture media was estimated to require a correction factor of approximately 1.44 for the 20x/0.8 NA Plan-Apochromat objectives according to previously established methods^56^. Therefore, the nominal (i.e., set in the software) Z-step size was corrected from 2 µm to the actual Z-step size of 2.88 µm and the nominal 0.5 µm Z-steps were corrected to 0.72 µm. After this adjustment the fluorescent sphere from optical control images became spherical instead of squat.

## 3. Quality control and dataset curation

### 3.1 Time-lapse annotations

Each movie was inspected and annotated by microscopists. Two microscopists assessed the cell health of each time-lapse and annotated if there was more cell death during the time-lapse than the baseline. When the two annotators disagreed a third one was consulted. The same method was used for assessing the presence of migrating cells. A time-lapse was annotated for the presence of migration, characterized by observing the majority of cells present in the first time point migrating by the 48 h time point. All the movies were also annotated manually for the presence of an extra colony or lumenoid at the time of migration onset (determined by the first time point showing the detachment of a healthy cell from the colony/lumenoid) as well as the presence of migrating cells coming from a colony or lumenoid outside of the field of view (FOV).

### 3.2 Analysis dataset curation

Only the first 48 h of the 60-hour movies were analyzed in this study. An additional 12 h were acquired to verify that the cells were not undergoing excessive cell death at the end of the acquisition, ensuring that cell health was optimal during the first 48 h of the movie. Time-lapses used for analysis were chosen based on the annotation criteria of positive cell health, presence of migration and no additional colony or lumenoids or migrating cells coming from a colony outside of the FOV were present in the FOV at the time of migration onset.

The resulting analysis dataset was comprised of 46 mEGFP-tagged H2B, 63 mEGFP-tagged Eomes, 35 mEGFP-tagged SOX-2, 54 mEGFP-tagged E-cadherin, and 70 mEGFP-tagged ZO-1 3D time-lapse movies with a total of 60 2D PLF colonies, 121 2D colonies, and 87 3D lumenoid conditions across cell lines.

## 4. Image analysis methods

### 4.1 Camera alignment matrix generation

The field of rings optical control images were acquired on the two cameras simultaneously, including bright-field and all fluorescence channels. To align these two cameras to each other we selected two fluorescent channels, one for each camera, with the closest wavelengths and segmented their field of rings pattern through a 2D spot filter. We matched rings between the two images through linear sum assignment optimization, weighing each potential match between nearby rings by their Euclidean distance (SciPy Python package). These matched points were used to estimate a similarity transformation matrix using scikit-image’s *estimate_transform()* function and adjusted the resulting transformation matrix to not produce any image scaling.

Downstream adjustment of images or coordinates based on alignment metrics from these optical controls was performed by applying the transformation matrix from the appropriate optical control image to the image/coordinates in question with scikit-image’s *warp()* and *warp_coords()* functions, respectively, and using nearest neighbor interpolation (scikit-image Python package).

### 4.2 All-cells mask generation from bright-field

To create colony masks representing the location of cells in the 3D images, called the “*all-cells masks*,” we utilized the 60 h time-lapse dataset of AAVS1mEGFP expressing cells imaged every 30 min. The mEGFP expressed from a safe harbor locus localizes to the cytoplasm and to a lesser extent the nucleus (cytoplasmic mEGFP), making it an ideal cell filling marker to define cell and therefore colony location. First, we used Otsu thresholding on the cytoplasmic mEGFP fluorescence images followed by a median filter of size 2×2×1 voxels to generate ground truth segmentation masks. Then, we trained a convolutional neural network (CNN)^57^ to predict the segmentation masks directly from the bright-field images from different conditions and different time points during the EMT process. We performed a multi-scale patch-based inference, where patches of different scales (P_i_, i = 1, 2,…, N; where N is the maximum number of patch scales, and each patch scale P_i_ has a different Z, Y, and X dimension) were used for model prediction. During all-cells mask generation, inference was performed at three different patch scales (N = 3, and P_1_= 16×128×128, P_2_ = 16×256×256, P_3_ = 16×512×512). The three probability maps (CNN output associated with each patch scale) generated during inference were then binarized using Otsu thresholding. Finally, by performing logical disjunction operation, the three binarized masks were merged to generate the all-cells mask. In total, we used 3,240 images for training and 1,440 for validation, with representative time points throughout the EMT process. The ground truth and predicted segmentation masks were validated by overlaying masks with the raw microscopy images and spot-checking approximately 15% of the data. After training, this model was used to predict binary colony masks for all 3D images analyzed in this manuscript.

### 4.3 Denoising method

Denoised images were processed using the Cellpose 3 denoise model to improve downstream segmentation or visualization of the low signal-to-noise ratio (SNR) inherent with fast live cell spinning disk confocal acquisition settings^58^. We found that the default Cellpose rescaling parameter applied a different rescaling factor to each Z-slice which improperly scaled out of focus plane pixel noise. To overcome this challenge, we determined an optimal constant pixel intensity scaling factor for each structure and applied it to the raw input images. These contrast-stretched images were then processed using the “denoise_cyto” image denoising model which enhances the raw image quality by reducing Poisson pixel noise which directly increases the SNR of denoised output stacks. Images of mEGFP-tagged Eomes and mEGFP-tagged H2B were denoised prior to downstream nuclear segmentation. Images of mEGFP-tagged ZO-1 taken every 3 min were denoised prior to hand annotation of the apical faces show in (Fig. 7D and Supplemental Movies 4-6).

### 4.4 3D nuclear instance segmentation

A robust 3D instance segmentation processing pipeline was developed for the H2B and Eomes mEGFP-tagged cell lines. Considerations were made through the workflow to overcome the limitations posed by large Z-step size and the presence of overlapping stacked nuclei. First, 3D image stacks were denoised as described above. Images were then segmented using the Cellpose cyto3 model, configured with a diameter of 60 pixels and a cell probability threshold of −2.0 to maximize segmentation accuracy and object detection sensitivity. To refine the 3D instance segmentation masks, a series of clipping and filtering steps were performed. First, Cellprofiler was used to filter low-intensity background artifacts and small cell debris from the segmentation masks by measuring object intensities and filtering objects based on a minimum integrated intensity threshold of 550. Next, a vectorized 3D clipping method was developed that applies an intensity threshold (set to 0.75) based on the mean pixel intensity within each object mask. Lastly, CellProfiler was employed for filtering any small objects that were created by the previous clipping step. This involves filtering objects based on integrated intensity, resulting in the final processed masks ready for downstream analysis. Nuclear segmentations and centroid locations were visualized by overlaying them with the raw microscopy images and manually checking approximately 50% of the dataset. Nuclear masks in upper Z-slices are impacted by spherical aberration and light scattering from refractive index mismatch and from cells in lower Z-slices, detrimentally impacting segmentations in upper Z-slices. Nuclear segmentations were validated for centroid location and used to determine migration timing, not for other applications. This comprehensive processing pipeline yields 3D segmented nuclei and their corresponding centroid locations with high detection sensitivity across the H2B and Eomes cell lines. The 3D segmented nuclei can be visualized using the web-based 3D Volume Viewer.

### 4.5 Basement membrane segmentation and 3D mesh generation

To observe cell migration during differentiation, we analyzed a cell’s location, given by its 3D nuclear instance segmentation centroid, relative to the lumenoid’s basement membrane surface (Supplemental Fig. S4: Mesh Generation). To do so, we first generated collagen IV segmentations using a dynamic UNet model^57^. To generate ground truths to train this model, we used an Otsu threshold on 21 collagen IV images (19 for training and two for validation) and manually curated the results to fill in small holes in the segmentations. The trained model was then used to predict the collagen IV probability maps for the rest of the image dataset containing 13,300 Z-stacks. To ensure that only the lumenoid in the center of the field of view was segmented, these probability maps were post-processed by thresholding the image at a probability of 0.25, determining the largest connected component, and masking out all other regions by setting their probabilities to 0. To generate 3D meshes from these segmentations, we set the threshold of the post-processed segmentation to 0.3 and randomly sampled the segmented regions to form a point cloud. The Z-coordinates of these points were adjusted so that the point cloud had isotropic dimensions. We then created an initial mesh in the shape of an inverted dome that was centered on the center of mass of the point cloud. The dome was created by generating an icosphere and removing the faces of its upper half. The height of this dome was 80% the height of the point cloud and its diameter was equal to the minimum width of the point cloud, as determined by its bounds in X, Y, and Z. We then used a Non Rigid Iterative Closest Points method to register the dome to the point cloud, generating a mesh that matched closely to the contours of the collagen IV basement membrane^59^. These meshes were validated by overlaying them, alongside the UNet segmentations, on top of the raw collagen IV signal for five randomly selected time points in each time-lapse processed.

Next, we used these collagen IV basement membrane meshes to determine whether a cell was inside or outside the lumenoid. First, the meshes were converted into watertight meshes, meaning that the open top left from cutting the initial icosphere in half was closed by adding additional faces over the top of the mesh^60^. Then the centroid of each segmented nucleus was used to determine if the cell was enclosed within the region of the membrane mesh. If the centroid was inside the mesh, the cell was considered inside the lumenoid; otherwise it was considered outside the lumenoid (Supplemental Fig. S4). Centroid coordinates were adjusted based on the alignment metrics determined from optical control images to correct for camera misalignment. Meshes were visually validated by comparing movies of approximately 25% of the generated meshes to their time-lapses of origin, showing both the basement membrane mesh and the centroids of nuclei color-coded by their classification.

## 5. Data analysis methods

### 5.1 Feature extraction from all-cells mask

The binary all-cells mask was used to extract the colony’s area at each Z-slice in addition to total intensity which was measured from the corresponding fluorescent channel. We aligned Z-positions from different time-lapses by subtracting the Z-slice closest to the glass from the absolute Z-slice, resulting in all colonies having a normalized Z at the glass of 0. The plane of the glass was determined by first summing the area present in each Z-plane of every all-cells mask across one entire time-lapse. Then, the first derivative of the sum of areas going from low to high Z was calculated, and the Z-slice where the derivative was maximal was set to have a normalized Z of 0 for that time-lapse.

### 5.2 Determination of migration timing using all-cells mask

Migration time for each movie was calculated by identifying the inflection point of the increase in colony area at the glass over time. The colony area at the glass was calculated by computing the area of the maximum intensity projection of the bottom two Z-slices of all-cells masks for each time point. The point of inflection was defined as the time point at which the second derivative of colony area at the glass was maximal between 17.5 and 40 h. The second derivative was computed by applying the Savitzky-Golay filter (SciPy Python library) to smooth the mean intensity values over the entire time-lapse and using a second order polynomial/quadratic polynomial over a window of 40 time points. The inflection point metric for migration timing was validated by comparing the error between multiple manual annotations of migration timing to the difference between manual and inflection point-based migration timings. We performed a non-parametric Kruskal-Wallis test followed by Mann-Whitney post-hoc test (using SciPy Stats and Scikit-posthoc Python libraries) to determine statistical significance of the difference between the migration timings for different conditions. Holm-Bonferroni method of p-value adjustment was performed for the post-hoc statistical analysis to adjust the p-values. p-value < .05 (alpha=0.05) was considered significant enough to reject the null hypothesis that the distributions underlying the two samples are the same.

### 5.3 Determination of migration timing using the basement membrane mesh

For 3D lumenoids, migration timing was also calculated by looking at the number of cells detected outside the basement membrane over time. For each time point in a movie, the number of nuclei outside the basement membrane was determined by classifying the centroid positions of the nuclei obtained from 3D instance segmentation to be either inside or outside the basement membrane. Migration time was then calculated by identifying the inflection point of the fraction of nuclei outside the basement membrane over time. This inflection point was computed by determining the time point at which the second derivative of the fraction of nuclei outside the basement membrane over time is maximum. The second derivative was computed by applying the Savitzky-Golay filter to smooth the mean intensity values for the time frame of 12-40 h using a second order polynomial/quadratic polynomial using a window of 30 time points.

### 5.4 Mean intensity over time

Mean intensity for each time point per movie was calculated by determining total intensity divided by total area over the first 10 Z-slices (starting at normalized z=0). To quantify transitions in variable mean intensity trajectories over time, different quantitative approaches were taken. SOX-2 mean-intensity trajectories are characterized by overall decay of intensity over time. To compare the dynamics of SOX-2 decay, we computed the time at which SOX-2 mean intensity decays to its half maximum (with respect to the minimum it reaches). The computation was performed by smoothing the mean intensity curve (Savitzky-Golay filter with a quadratic polynomial) over a window of 10 time points and calculating the time point closest to the mid-point between maximum and minimum intensity of the SOX-2 trajectory.

EOMES mean-intensity trajectories increase then decrease over the course of imaging. To compute the time corresponding to peak EOMES intensity, we first smoothed the mean intensity trajectory (Savitzky-Golay filter with a quadratic polynomial) over a window of 10 time points to factor out noise. We then computed the maximum intensity of the trajectory and the corresponding time point. E-cadherin mean intensity trajectories are characterized by a period of high signal followed by decay. To compute the inflection time of decay in E-cadherin, the second derivative of mean intensity over time was calculated using the Savitzky-Golay filter with a quadratic polynomial (window=40). The inflection time was determined by identifying the time point of the minimum of the second derivative between 17.5 and 39 h. To statistically compare the quantitative metrics computed from mean intensity trajectories between the three experimental conditions, we first performed a non-parametric Kruskal-Wallis test followed by Mann-Whitney post-hoc test (using SciPy Stats and Scikit-posthoc Python libraries). For the Mann-Whitney post-hoc test, we used Holm-Bonferroni p-value correction to compute pair-wise p-values.

### 5.5 Characterization of apical dynamics using ZO-1

To characterize the change in ZO-1 during migration, the mean intensity inside the all-cells mask was measured at each Z-slice at each time point and the mean intensity, Z-height, and time were plotted together as a heatmap (with a lower threshold of 50,000 pixels per Z-slice). To visualize apical closure over time, a maximum Z-projection of two slices of denoised ZO-1 images was generated, visualized in pseudo-color, and apical faces were manually outlined.

## 6. Visualization

### 6.1 3D Volume Viewer

All OME-Zarr datasets associated with this paper can be accessed and viewed without downloading through the web-based 3D Volume Viewer available at http://volumeviewer.allencell.org/. This viewer is designed to optimize performance and speed, with capabilities to automatically select the appropriate resolution while rendering interactive 3D time series. Additionally, the viewer supports the automatic combination of different image files, enabling simultaneous display and overlay of raw microscopy images and their corresponding segmentations, even when these images are stored in separate files. See Supplemental Table 1 for 3D Volume Viewer links for each time-lapse shown in each figure.

### 6.2 AGAVE rendering

3D volumetric renderings were generated using AGAVE. AGAVE uses a physically based lighting and rendering algorithm to produce high quality views of volumetric data. For more info about AGAVE, visit https://www.allencell.org/pathtrace-rendering.html and https://github.com/allen-cell-animated/agave.

### 6.3 BioFile Finder

The BioFile Finder (BFF) is an open-use web application created for easy access, collaboration, and sharing of datasets through rich metadata search, filter, sort, and direct viewing in common industry applications or in our web-based 3D Volume Viewer (https://biofile-finder.allencell.org). Links to data released with this manuscript in BFF are provided in Methods section 7.

## 7. Data availability

We release all time-lapse data used in this study in the OME-Zarr format to democratize their access. Multi-scene czi files were split into mono-scene ome.tiff files using Python packages bioio and bioio_czi. Multi-scene ND2 movies were split into mono-scene ND2 files using NIS ELEMENT VIEWER 5.21. The resulting mono-scene movies were then converted to OME-Zarr. During this lossless process, channels were renamed for consistency and Z-pixel size was updated to reflect the actual pixel size instead of the nominal one (see Methods section 2.4).

The resulting OME-Zarr movies and corresponding segmentations are made available for download on Quilt and easy online viewing using BioFile Finder (BFF) and 3D Volume Viewer web applications. Users can programmatically access and download the data at the following Quilt repository: (https://open.quiltdata.com/b/allencell/tree/aics/emt_timelapse_dataset/). A complete description of dataset annotations is available in the manifest: Imaging_and_segmentation_data_column_definition.csv. All features extracted from the different segmentations are provided in the manifest Image_analysis_extracted_features.csv. A complete description of all columns of this file is provided in the manifest Image_analysis_extracted_features_column_description.csv. Similarly, all features extracted for the nuclear localization inside/outside of basement membrane analysis are provided in Migration_timing_through_mesh_extracted_features.csv, accompanied by a description of each column in Migration_timing_through_mesh_extracted_features_column_description.csv. Annotations describing the microscope settings during acquisition and biological description of the sample imaged are also provided in the manifest: imaging_and_segmentation_data.csv.

BFF enables users to dynamically search and visualize all movies and corresponding segmentations utilized in this study without downloading them. All time-lapses can be visualized by selecting the movie of interest, clicking the Open File button, and selecting the desired visualization option from the drop-down menu. To facilitate access, we provide two primary BFF links: the first link organizes the data by biological conditions and gene, while the second link organizes the data according to their application in the study. Users can organize files in various combinations based on metadata to suit their specific needs. We encourage users to customize their queries dynamically to best fit their research requirements with 53 different annotation criteria to choose from.

Data organized in BFF by biological condition and mEGFP-tagged gene: https://biofile-finder.allencell.org/app?group=Experimental+Condition&group=Gene&source=%7B%22name%22%3A%22imaging_and_segmentation_data.csv+%288%2F15%2F2024+4%3A26%3A03+PM%29%22%2C%22type%22%3A%22csv%22%2C%22uri%22%3A%22https%3A%2F%2Fallencell.s3.amazonaws.com%2Faics%2Femt_timelapse_dataset%2Fmanifests%2Fimaging_and_segmentation_data.csv%3FversionId%3DWmTjARBNL4rNJhV4N7YFYr2dKHWlCHwc%22%7D&sourceMetadata=%7B%22name%22%3A%22Imaging_and_segmentation_data_column_description.csv+%288%2F15%2F2024+4%3A26%3A02+PM%29%22%2C%22type%22%3A%22csv%22%2C%22uri%22%3A%22https%3A%2F%2Fallencell.s3.amazonaws.com%2Faics%2Femt_timelapse_dataset%2Fmanifests%2FImaging_and_segmentation_data_column_description.csv%3FversionId%3D.bmbr.UUT06F9nupeuwxVBYuTMyKyYu6%22%7D

Data pre-organized in BFF by use in this manuscript: https://biofile-finder.allencell.org/app?group=Used+For&group=Experimental+Condition&group=Gene&source=%7B%22name%22%3A%22imaging_and_segmentation_data.csv+%288%2F15%2F2024+4%3A26%3A03+PM%29%22%2C%22type%22%3A%22csv%22%2C%22uri%22%3A%22https%3A%2F%2Fallencell.s3.amazonaws.com%2Faics%2Femt_timelapse_dataset%2Fmanifests%2Fimaging_and_segmentation_data.csv%3FversionId%3DWmTjARBNL4rNJhV4N7YFYr2dKHWlCHwc%22%7D&sourceMetadata=%7B%22name%22%3A%22Imaging_and_segmentation_data_column_description.csv+%288%2F15%2F2024+4%3A26%3A02+PM%29%22%2C%22type%22%3A%22csv%22%2C%22uri%22%3A%22https%3A%2F%2Fallencell.s3.amazonaws.com%2Faics%2Femt_timelapse_dataset%2Fmanifests%2FImaging_and_segmentation_data_column_description.csv%3FversionId%3D.bmbr.UUT06F9nupeuwxVBYuTMyKyYu6%22%7D

## 8. Code availability

Custom written code was central to the analysis and conclusions of this paper. All necessary code for image analysis and data analysis has been shared publicly on GitHub at https://github.com/AllenCell/EMT_image_analysis and https://github.com/AllenCell/EMT_data_analysis.

