## Supplementary Materials for "A human induced pluripotent stem (hiPS) cell model for the holistic study of epithelial to mesenchymal transitions (EMTs)"

**Supplemental information in this document includes Figures S1-S6, Captions of Supplemental Movies 1-6, and Supplemental Table 1.**

**Other Supplemental Materials for this manuscript include Movies 1-6**

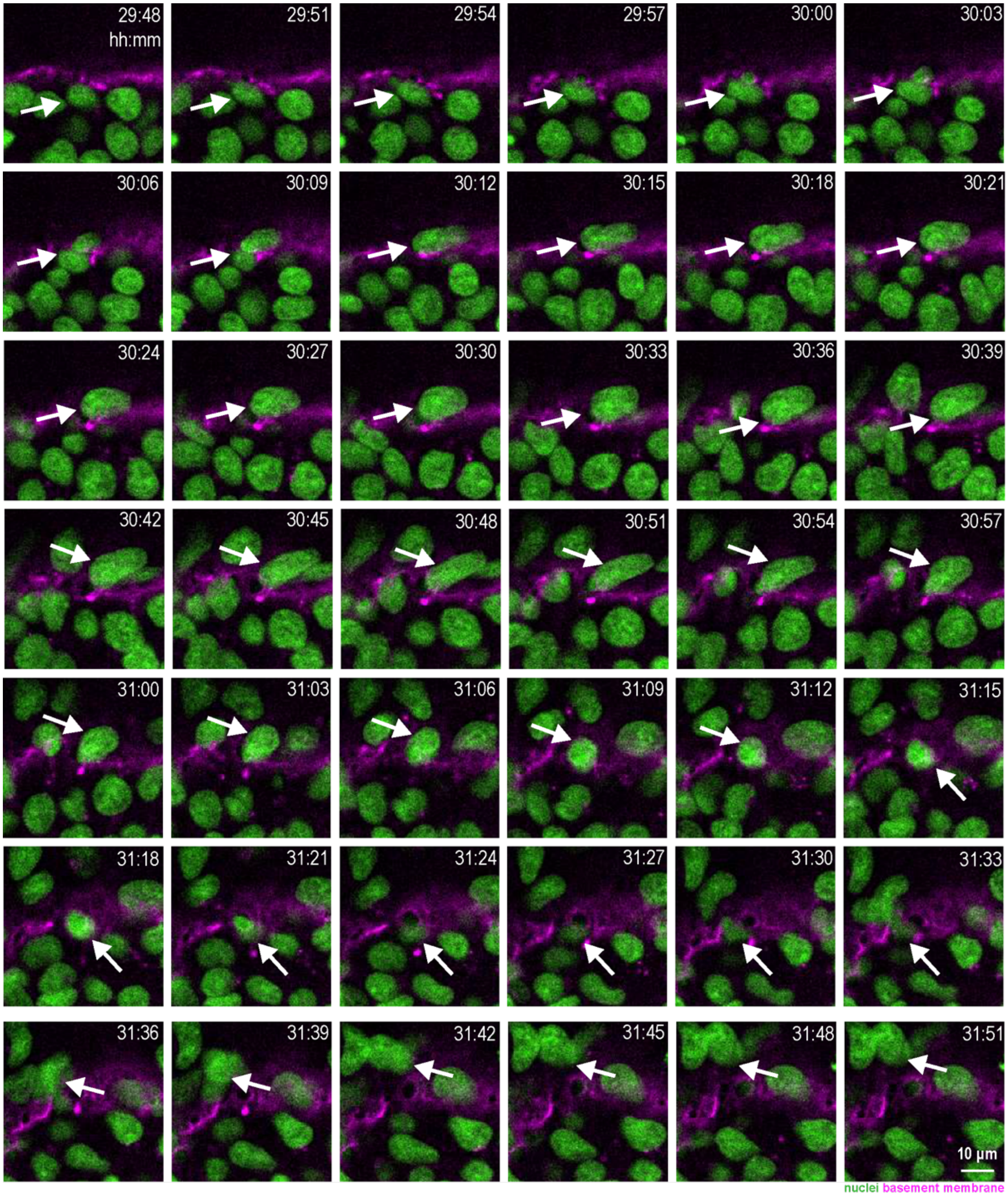


**Fig. S1: Exiting the lumenoid and the basement membrane is a reversible process.** Three-minute interval time-lapse images of a *3D lumenoid EMT* condition showing the nucleus (mEGFP-tagged H2B; green; see arrow) of a single cell repeatedly crossing the basement membrane (human-specific anti-collagen IV antibody; magenta). Images are single Z-slices at the approximate middle, shown in XY. Timestamps indicate the time post-EMT induction. Scale bar: 10 µm (see Supplemental Movie 3).

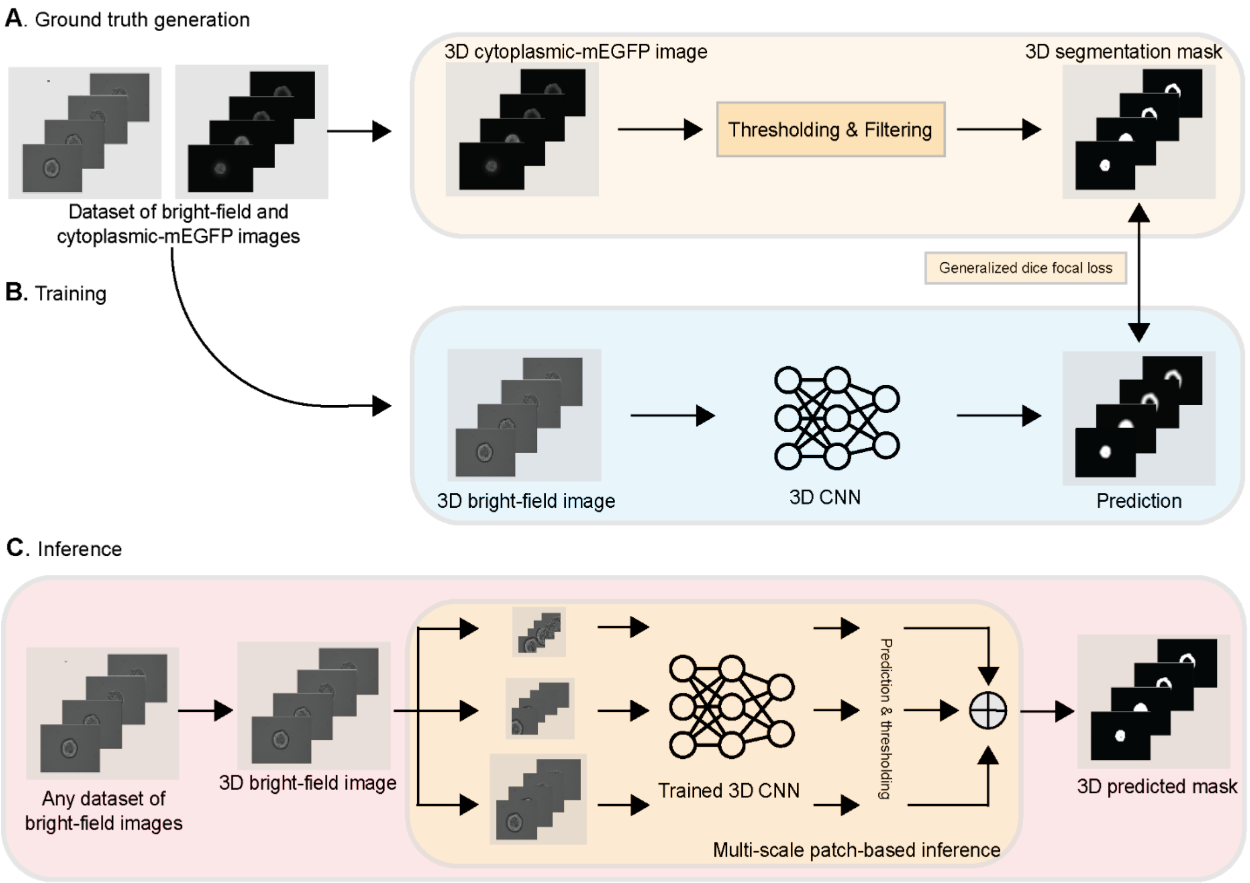


**Fig. S2: A deep learning model for predicting all-cells masks in 3D from bright-field Z-stacks using cytoplasmic mEGFP expressing hiPS cells for ground truth generation.** **A.** To generate ground truth masks, Otsu thresholding and median filtering of 3D cytoplasmic mEGFP images was performed. **B.** A 3D CNN model was trained using these 3D segmentation masks (for the generalized dice focal loss calculation) and their respective 3D bright-field images. **C.** During inference, model predictions were generated for each 3D bright-field Z-stack using a multi-scale patch-based approach. Bright-field-generated masks were produced by merging the Otsu thresholded model predictions associated with each patch scale. Overlays of the resulting all-cells masks with raw bright-field images were generated for visual validation (see Methods 4.2).

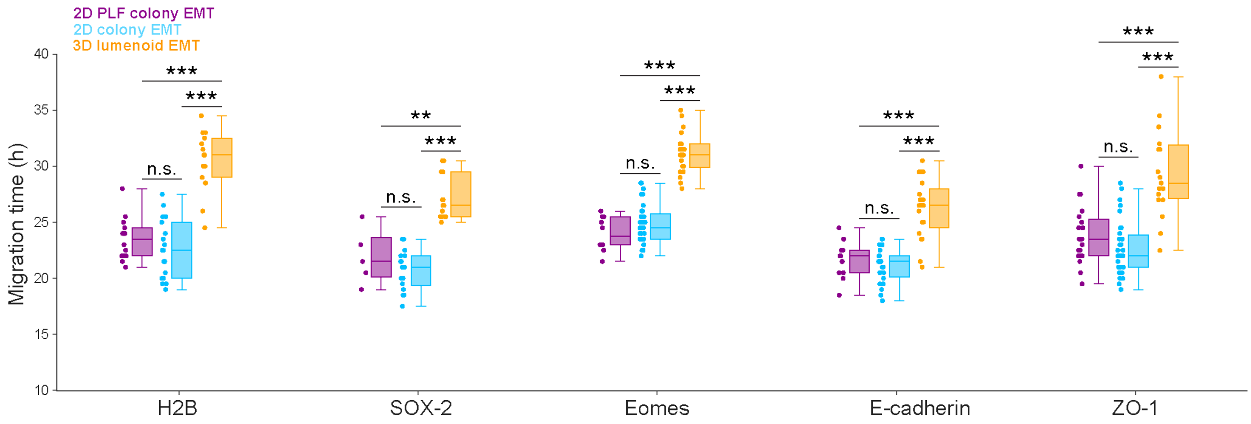


**Fig. S3: Reproducible differences in 2D and 3D EMT conditions exist despite small differences in timing between cell lines** Estimated time of migration determined from the all-cells masks as shown in Fig. 5 A-D and described in Methods section 5.2. Statistical significance results after Holm-Bonferroni p-value adjustment for each cell line for the following paired conditions (*2D PLF colony EMT* vs. *2D colony EMT*, *2D PLF colony EMT* vs. *3D lumenoid EMT*, *2D colony EMT* vs. *3D lumenoid EMT*) were as follows: H2B p=(.30, $3.7\times10^{-5}$, $2.0\times10^{-5}$), SOX-2 p=(.41, $6.8\times10^{-3}$, $1.2\times10^{-5}$), Eomes p=(.21, $1.9\times10^{-5}$, $4.4\times10^{-9}$), E-cadherin p=(.36, $2.8\times10^{-4}$, $2\times10^{-6}$), and ZO-1 p=(.086, $4.9\times10^{-5}$, $8.5\times10^{-7}$). Statistically significant p-values <.001 were designated with ***, p-values .001 to .01 were designated with **, and p-value >=.05 was considered not significant (n.s.). The sample sizes from each cell line for each condition (*2D PLF colony EMT*, *2D colony EMT*, *3D lumenoid EMT*) were as follows: H2B n=(14, 18, 14), Eomes n=(10, 32, 21), E-cadherin n=(11, 23, 20), and ZO-1 n=(20, 31, 19).

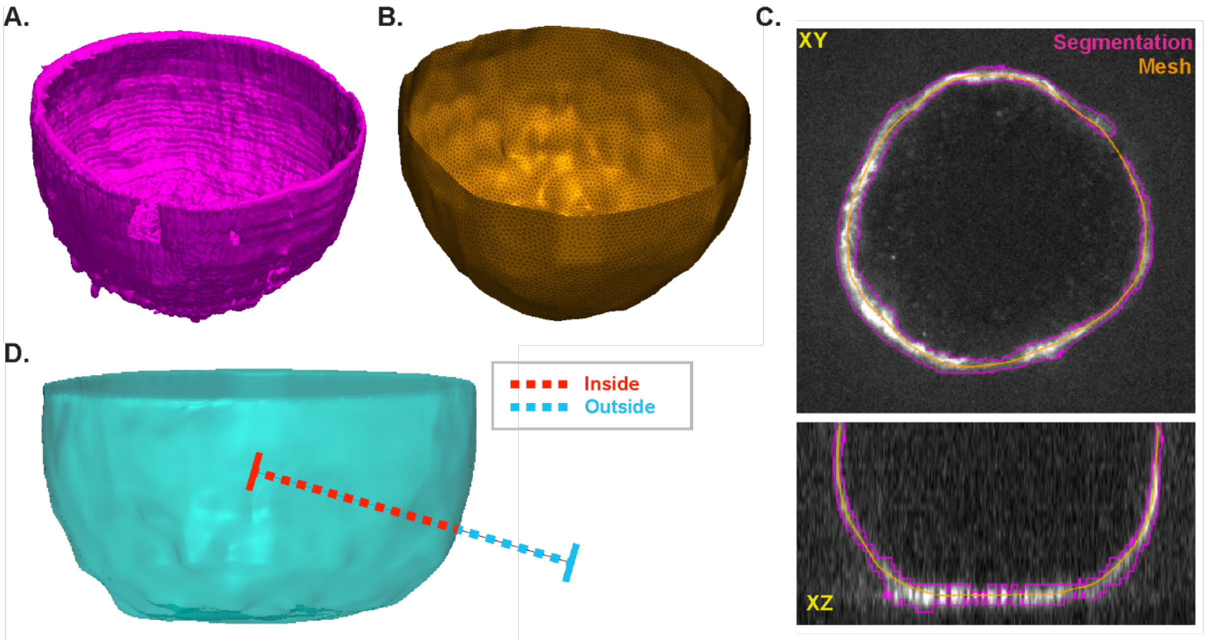


**Fig. S4: Basement membrane mesh generation and inside/outside classification of cells. A.** The basement membrane is segmented using a UNet model (shown here as a 3D voxel object; magenta). **B.** The segmented region in A is randomly sampled to generate a point cloud and a 3D mesh of an inverted dome is registered to this point cloud using a Non Rigid Iterative Closest Points method (orange). **C.** Overlay of the UNet segmentation in A (magenta) and the registered mesh in B (orange) over the original collagen IV signal (white). **D.** The registered 3D mesh is converted to a watertight object (the open top is closed; cyan) and is used to determine if a nucleus is inside or outside of the lumenoid.

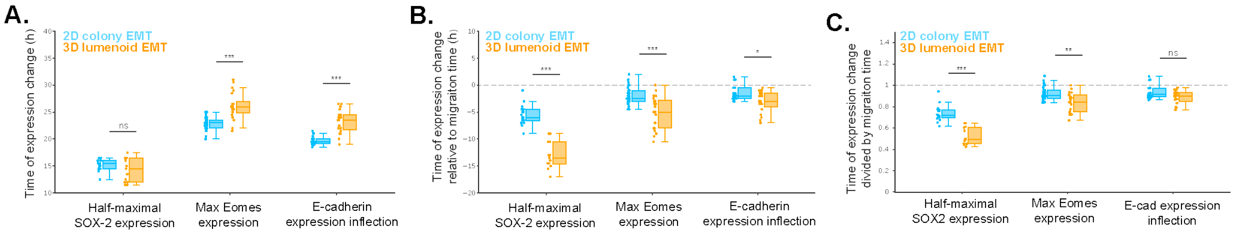


**Fig. S5 Relating the timing of changes in marker expression to the time of EMT induction or the start of cell migration A.** Timing of expression change relative to induction of EMT (time 0) in h. **B**. Timing of expression change relative to migration time in h, with negative numbers representing time before migration. **C**. Timing of expression change normalized between the time of induction and migration, with 0 being time of EMT induction and 1 being time of migration. Statistically significant p-values <.001 were designated with ***, p-values .001 to .01 with **, p-values .01 to .05 with *, and p-value >=.05 was considered not significant (n.s.).

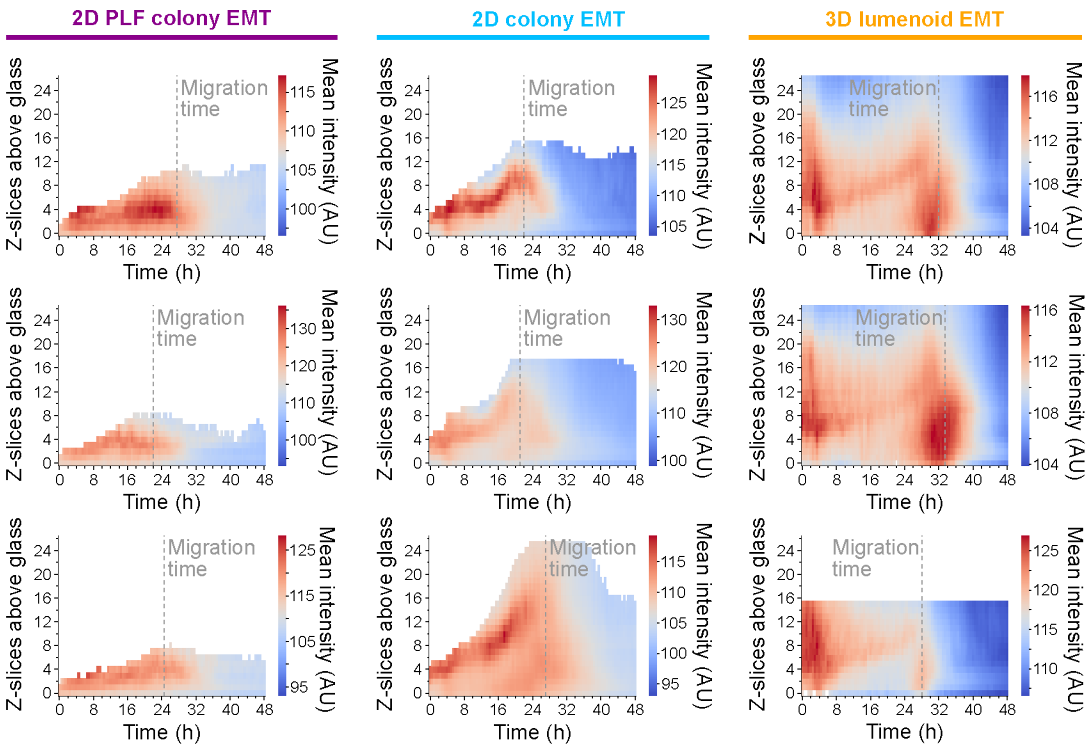


**Fig. S6. The pattern of ZO-1 expression and localization is reproducible across multiple examples from each EMT condition** Three additional representative heatmaps for each EMT condition showing the mean fluorescence intensity of ZO-1 as a function of height above the glass over time as measured using the all-cells masks. The migration time for each example is indicated on the plot with a gray dashed vertical line.

**Supplemental Movies Captions:**

**Supplemental Movie 1. Time-lapse corresponding to the images in Fig. 4C, showing an example cell leaving the lumenoid by deforming to fit through a narrow hole in the basement membrane at three-minute time resolution.** mEGFP-tagged H2B was used to visualize the nucleus (green) and human-specific anti-collagen IV antibody was used to visualize the basement membrane (magenta) in *3D Lumenoid EMT*. The movie is a single Z-slice and starts 29:57 (hh:mm) after EMT induction. Scale bar is 10 µm and frame rate is 2 fps.

**Supplemental Movie 2: Time-lapse corresponding to the images in Fig. 4D, showing an example cell leaving the lumenoid by passing, undeformed, through larger holes in the basement membrane at a three-minute interval.** mEGFP-tagged H2B was used to visualize the nucleus (green) and human-specific anti-collagen IV antibody was used to visualize the basement membrane (magenta) in *3D Lumenoid EMT*. The movie is a single Z-slice and starts 33:03 (hh:mm) after EMT induction. Scale bar is 10 µm and frame rate is 3 fps.

**Supplemental Movie 3: Time-lapse corresponding to the images in Fig. S1, showing that exiting the lumenoid through the basement membrane is a reversible process at three-minute time resolution.** mEGFP-tagged H2B was used to visualize the nucleus (green) and human-specific anti-collagen IV antibody was used to visualize the basement membrane (magenta) in *3D Lumenoid EMT*. The white arrow points to a single nucleus repeatedly crossing the basement membrane. The movie is a single Z-slice and starts 29:48 (hh:mm) after EMT induction. Scale bar is 10 µm and frame rate is 4 fps.

**Supplemental Movie 4. Time-lapse corresponding to images in Fig. 7D, showing a *2D PLF Colony EMT* example at three-minute time resolution to visualize apical closure.** mEGFP-tagged ZO-1 was used to mark tight junctions and apical faces. The movie is a maximum intensity projection of two Z-slices of denoised fluorescence data shown in pseudo-color and starts 20:48 (hh:mm) after EMT induction. Scale bar is 10 µm and frame rate is 3 fps.

**Supplemental Movie 5. Time-lapse corresponding to images in Fig. 7D, showing a *2D Colony EMT* example at three-minute time resolution to visualize apical closure.** mEGFP-tagged ZO-1 was used to mark tight junctions and apical faces. The movie is a maximum intensity projection of two Z-slices of denoised fluorescence data shown in pseudo-color and starts at 20:18 (hh:mm) after EMT induction. Scale bar is 10 µm and frame rate is 3 fps.

**Supplemental Movie 6. Time-lapse corresponding to the images in Fig. 7D, showing a *3D Lumenoid EMT* example at three-minute time resolution to visualize apical closure.** mEGFP-tagged ZO-1 was used to mark tight junctions and apical faces. The movie is a maximum intensity projection of two Z-slices of denoised fluorescence data shown in pseudo-color and starts at 21:00 (hh:mm) after EMT induction. Scale bar is 10 μm and frame rate is 3 fps.

**Supplemental Table 1: Web-based 3D Volume Viewer links for each time-lapse shown in figures.**

| **Figure 2: Environmental Matrigel changes hiPS cell colony geometry during growth.** | | | |
|---|---|---|---|
| *Section* | *EMT condition* | *Structure* | *3D Volume Viewer link* |
| Fig. 2A | hiPS cell 2D colony growth | mEGFP-tagged H2B, human-specific anti-collagen IV antibody | https://volumeviewer.allencell.org/viewer?url=https%3A%2F%2Fallencell.s3.amazonaws.com%2Faics%2Femt_timelapse_dataset%2Fdata%2F3500005824_32_raw_converted.ome.zarr&mode=maxproject&c0=cps:0:0:1:78:0:1:89:1:1:255:1:1,rmp:78:89&c1=ven:1,col:6fba11,cps:0:0:1:6:0:1:21:1:1:255:1:1,rmp:3:71&c2=ven:1,col:bd10e0,cps:0:0:1:17:0:1:27:1:1:255:1:1,rmp:17:55 |
| Fig. 2B | hiPS cell 2D to 3D lumenoid formation | mEGFP-tagged H2B, human-specific anti-collagen IV antibody | https://volumeviewer.allencell.org/viewer?url=https%3A%2F%2Fallencell.s3.amazonaws.com%2Faics%2Femt_timelapse_dataset%2Fdata%2F3500005824_16_raw_converted.ome.zarr&mode=maxproject&c0=cps:0:0:1:83:0:1:96:1:1:255:1:1,rmp:83:96&c1=ven:1,col:6fba11,cps:0:0:1:9:0:1:36:1:1:255:1:1,rmp:6:86&c2=ven:1,col:bd10e0,cps:0:0:1:60:0:1:98:1:1:255:1:1,rmp:56:253 |
| **Figure 3: EMT is influenced by starting hiPS cell colony geometry, environmental Matrigel, and substrate adhesiveness.** | | | |
| *Section* | *EMT condition* | *Structure* | *3D Volume Viewer link* |
| Fig. 3A | 2D colony no Matrigel EMT | mEGFP-tagged H2B, human-specific anti-collagen IV antibody | https://volumeviewer.allencell.org/viewer?url=https%3A%2F%2Fallencell.s3.amazonaws.com%2Faics%2Femt_timelapse_dataset%2Fdata%2F3500006350_48_raw_converted.ome.zarr&mode=maxproject&c0=cps:0:0:1:113:0:1:131:1:1:255:1:1,rmp:113:131&c1=ven:1,col:6fba11,cps:0:0:1:7:0:1:28:1:1:255:1:1,rmp:0:69&c2=ven:1,col:bd10e0,cps:0:0:1:87:0:1:136:1:1:255:1:1,rmp:65.243:332.78 |
| Fig. 3B | 2D colony EMT | mEGFP-tagged H2B, human-specific anti-collagen IV antibody | https://volumeviewer.allencell.org/viewer?url=https%3A%2F%2Fallencell.s3.amazonaws.com%2Faics%2Femt_timelapse_dataset%2Fdata%2F3500006256_12_raw_converted.ome.zarr&mode=maxproject&c0=cps:0:0:1:89:0:1:106:1:1:255:1:1,rmp:89:106&c1=ven:1,col:6fba11,cps:0:0:1:10:0:1:29:1:1:255:1:1,rmp:6:93.043&c2=ven:1,col:bd10e0,cps:0:0:1:102:0:1:158:1:1:255:1:1,rmp:112.35:331.19 |
| Fig. 3C | 2D PLF colony EMT | mEGFP-tagged H2B, human-specific anti-collagen IV antibody | https://volumeviewer.allencell.org/viewer?url=https%3A%2F%2Fallencell.s3.amazonaws.com%2Faics%2Femt_timelapse_dataset%2Fdata%2F3500005828_30_raw_converted.ome.zarr&mode=maxproject&c0=cps:0:0:1:117:0:1:134:1:1:255:1:1,rmp:117:134&c1=ven:1,col:6fba11,cps:0:0:1:13:0:1:28:1:1:255:1:1,rmp:13:110&c2=ven:1,col:bd10e0,cps:0:0:1:34:0:1:52:1:1:255:1:1,rmp:21.476:149.65 |
| Fig. 3D | 3D lumenoid EMT | mEGFP-tagged H2B, human- | https://volumeviewer.allencell.org/viewer?url=https%3A%2F%2Fallencell.s3.amazonaws.com%2Faics%2Femt_timelapse_dataset%2Fdata%2F3500005824_36_raw_converted.ome.zarr&mode=maxproject&c0=c |

| | | | |
|---|---|---|---|
| | | specific anti-collagen IV antibody | ps:0:0:1:101:0:1:129:1:1:255:1:1,rmp:101:129&c1=ven:1,col:6fba11,cps:0:0:1:13:0:1:51:1:1:255:1:1,rmp:13:130.12&c2=ven:1,col:bd10e0,cps:0:0:1:33:0:1:55:1:1:255:1:1,rmp:32.131:156.55 |
| **Figure 4: Cells can leave lumenoids by deforming to fit through narrow holes in the basement membrane or by passing, undeformed, through larger holes.** | | | |
| *Section* | *EMT condition* | *Structure* | *3D Volume Viewer link* |
| Fig. 4A | 3D lumenoid EMT | mEGFP-tagged H2B, human-specific anti-collagen IV antibody | https://volumeviewer.allencell.org/viewer?url=https%3A%2F%2Fallencell.s3.amazonaws.com%2Faics%2Femt_timelapse_dataset%2Fdata%2F3500005824_36_raw_converted.ome.zarr&c0=cps:0:0:1:101:0:1:129:1:1:255:1:1,rmp:101:129&c1=ven:1,col:6fba11,cps:0:0:1:13:0:1:51:1:1:255:1:1,rmp:13:130&c2=ven:1,col:bd10e0,cps:0:0:1:33:0:1:55:1:1:255:1:1,rmp:48:54.008 |
| Fig. 4B | 3D lumenoid EMT | human-specific anti-collagen IV antibody | https://volumeviewer.allencell.org/viewer?url=https%3A%2F%2Fallencell.s3.amazonaws.com%2Faics%2Femt_timelapse_dataset%2Fdata%2F3500005824_36_raw_converted.ome.zarr&mode=maxproject&c0=cps:0:0:1:101:0:1:129:1:1:255:1:1,rmp:101:129&c1=col:6fba11,cps:0:0:1:13:0:1:51:1:1:255:1:1,rmp:13:130&c2=ven:1,col:ffffff,cps:0:0:1:33:0:1:55:1:1:255:1:1,rmp:13:262.06 |
| Fig. 4C | 3D lumenoid EMT | mEGFP-tagged H2B, human-specific anti-collagen IV antibody | see Supplemental Movie 1 |
| Fig. 4D | 3D lumenoid EMT | mEGFP-tagged H2B, human-specific anti-collagen IV antibody | see Supplemental Movie 2 |
| **Figure 5: Migration timing differs for 2D colony and 3D lumenoid geometries as determined by all-cells mask-derived systematic measurements of colony spreading** | | | |
| *Section* | *EMT condition* | *Structure* | *3D Volume Viewer link* |
| Fig. 5A | 2D colony EMT | N/A (bright-field) | https://volumeviewer.allencell.org/viewer?url=https%3A%2F%2Fallencell.s3.amazonaws.com%2Faics%2Femt_timelapse_dataset%2Fdata%2F3500005551_8_raw_converted.ome.zarr&mode=maxproject&reg=0:1,0:1,0.467:0.5&c0=ven:1,col:ffffff,cps:0:0:1:98:0:1:114:1:1:255:1:1,rmp:3.4182:166.13&c1=col:6fba11,cps:0:0:1:120:0:1:157:1:1:255:1:1,rmp:120:157&c2=col:bd10e0,cps:-10.303:0:1:79.848:0:1:129.13:1:1:296.21:1:1,rmp:79.848:129.13 |

| | | | |
|---|---|---|---|
| Fig. 5A | 3D lumenoid EMT | N/A (bright-field) | https://volumeviewer.allencell.org/viewer?url=https%3A%2F%2Fallencell.s3.amazonaws.com%2Faics%2Femt_timelapse_dataset%2Fdata%2F3500005551_46_raw_converted.ome.zarr&mode=maxproject&reg=0:1,0:1,0.467:0.5&c0=ven:1,col:ffffff,cps:0:0:1:100:0:1:131:1:1:255:1:1,rmp:0:165.67&c1=col:6fba11,cps:0:0:1:12:0:1:15:1:1:255:1:1,rmp:12:15&c2=col:bd10e0,cps:0:0:1:42:0:1:69:1:1:255:1:1,rmp:42:69 |
| **Figure 6: Fluorescence in bright-field-generated all-cells masks of EMT markers, but not SOX-2 show similar qualitative dynamics to that of migration timing.** | | | |
| *Section* | *EMT condition* | *Structure* | *3D Volume Viewer link* |
| Fig. 6A | 2D PLF colony EMT | mEGFP-tagged SOX-2 | https://volumeviewer.allencell.org/viewer?url=https%3A%2F%2Fallencell.s3.amazonaws.com%2Faics%2Femt_timelapse_dataset%2Fdata%2F3500005827_27_raw_converted.ome.zarr&mode=maxproject&c0=cps:0:0:1:88:0:1:99:1:1:255:1:1,rmp:88:99&c1=ven:1,col:ffffff,cps:0:0:1:57:0:1:77:1:1:255:1:1,rmp:50:130&c2=col:bd10e0,cps:0:0:1:29:0:1:46:1:1:255:1:1,rmp:14.357:209.42 |
| Fig. 6A | 2D colony EMT | mEGFP-tagged SOX-2 | https://volumeviewer.allencell.org/viewer?url=https%3A%2F%2Fallencell.s3.amazonaws.com%2Faics%2Femt_timelapse_dataset%2Fdata%2F3500005827_33_raw_converted.ome.zarr&mode=maxproject&c0=cps:0:0:1:122:0:1:139:1:1:255:1:1&c1=ven:1,col:ffffff,cps:0:0:1:51:0:1:102:1:1:255:1:1,rmp:39.424:130.95&c2=col:bd10e0,cps:0:0:1:77:0:1:124:1:1:255:1:1,rmp:77:124 |
| Fig. 6A | 3D lumenoid EMT | mEGFP-tagged SOX-2 | https://volumeviewer.allencell.org/viewer?url=https%3A%2F%2Fallencell.s3.amazonaws.com%2Faics%2Femt_timelapse_dataset%2Fdata%2F3500005827_45_raw_converted.ome.zarr&mode=maxproject&c0=cps:0:0:1:82:0:1:114:1:1:255:1:1,rmp:82:114&c1=ven:1,col:ffffff,cps:0:0:1:49:0:1:87:1:1:255:1:1,rmp:33:133.55&c2=col:bd10e0,cps:0:0:1:44:0:1:73:1:1:255:1:1,rmp:44:73 |
| Fig. 6E | 2D PLF colony EMT | mEGFP-tagged Eomes | https://volumeviewer.allencell.org/viewer?url=https%3A%2F%2Fallencell.s3.amazonaws.com%2Faics%2Femt_timelapse_dataset%2Fdata%2F3500005551_3_raw_converted.ome.zarr&mode=maxproject&c0=cps:0:0:1:104:0:1:118:1:1:255:1:1,rmp:104:118&c1=ven:1,col:ffffff,cps:0:0:1:128:0:1:165:1:1:255:1:1,rmp:78.907:355.61&c2=col:bd10e0,cps:0:0:1:36:0:1:56:1:1:255:1:1,rmp:36:56 |
| Fig. 6E | 2D colony EMT | mEGFP-tagged Eomes | https://volumeviewer.allencell.org/viewer?url=https%3A%2F%2Fallencell.s3.amazonaws.com%2Faics%2Femt_timelapse_dataset%2Fdata%2F3500005551_9_raw_converted.ome.zarr&mode=maxproject&c0=cps:0:0:1:98:0:1:114:1:1:255:1:1,rmp:98:114&c1=ven:1,col:ffffff,cps:0:0:1:112:0:1:152:1:1:255:1:1,rmp:68.189:371.1&c2=col:bd10e0,cps:0:0:1:6:0:1:10:1:1:255:1:1,rmp:6:10 |
| Fig. 6E | 3D lumenoid EMT | mEGFP-tagged Eomes | https://volumeviewer.allencell.org/viewer?url=https%3A%2F%2Fallencell.s3.amazonaws.com%2Faics%2Femt_timelapse_dataset%2Fdata%2F3500005551_46_raw_converted.ome.zarr&mode=maxproject&c0=cps:0:0:1:100:0:1:131:1:1:255:1:1,rmp:100:131&c1=ven:1,col:ffffff,cps:0:0:1:12:0:1:15:1:1:255:1:1,rmp:7.6465:35.303&c2=col:bd10e0,cps:0:0:1:42:0:1:69:1:1:255:1:1,rmp:42:69 |

| | | | |
|---|---|---|---|
| Fig. 6I | 2D PLF colony EMT | mEGFP-tagged E-cadherin | https://volumeviewer.allencell.org/viewer?url=https%3A%2F%2Fallencell.s3.amazonaws.com%2Faics%2Femt_timelapse_dataset%2Fdata%2F3500006212_41_raw_converted.ome.zarr&mode=maxproject&c0=cps:0:0:1:99:0:1:111:1:1:255:1:1,rmp:99:111&c1=ven:1,col:ffffff,cps:0:0:1:42:0:1:60:1:1:255:1:1,rmp:34.084:100.95&c2=col:bd10e0,cps:0:0:1:61:0:1:96:1:1:255:1:1,rmp:61:96 |
| Fig. 6I | 2D colony EMT | mEGFP-tagged E-cadherin | https://volumeviewer.allencell.org/viewer?url=https%3A%2F%2Fallencell.s3.amazonaws.com%2Faics%2Femt_timelapse_dataset%2Fdata%2F3500006212_15_raw_converted.ome.zarr&mode=maxproject&slice=0.5,0.5,0.467&c0=cps:0:0:1:134:0:1:153:1:1:255:1:1,rmp:134:153&c1=ven:1,col:ffffff,cps:0:0:1:28:0:1:41:1:1:255:1:1,rmp:19.37:73.606&c2=col:bd10e0,cps:0:0:1:81:0:1:129:1:1:255:1:1,rmp:81:129 |
| Fig. 6I | 3D lumenoid EMT | mEGFP-tagged E-cadherin | https://volumeviewer.allencell.org/viewer?url=https%3A%2F%2Fallencell.s3.amazonaws.com%2Faics%2Femt_timelapse_dataset%2Fdata%2F3500006079_22_raw_converted.ome.zarr&mode=maxproject&c0=cps:0:0:1:100:0:1:119:1:1:255:1:1,rmp:100:119&c1=ven:1,col:ffffff,cps:0:0:1:69:0:1:95:1:1:255:1:1,rmp:64.491:139.52&c2=col:bd10e0,cps:0:0:1:65:0:1:103:1:1:255:1:1,rmp:65:103 |
| **Figure 7: hiPS cells undergoing EMT constrict their apical faces to a point before delaminating in all hiPS cell-EMT conditions.** | | | |
| *Section* | *EMT condition* | *Structure* | *3D Volume Viewer link* |
| Fig. 7A | 2D PLF colony EMT | mEGFP-tagged ZO-1 | https://volumeviewer.allencell.org/viewer?url=https%3A%2F%2Fallencell.s3.amazonaws.com%2Faics%2Femt_timelapse_dataset%2Fdata%2F3500005829_4_raw_converted.ome.zarr&mode=maxproject&c0=cps:0:0:1:108:0:1:127:1:1:255:1:1,rmp:108:127&c1=ven:1,col:ffffff,cps:0:0:1:30:0:1:42:1:1:255:1:1,rmp:30:65.495&c2=col:bd10e0,cps:0:0:1:13:0:1:21:1:1:255:1:1,rmp:13:21 |
| Fig. 7A | 2D colony EMT | mEGFP-tagged ZO-1 | https://volumeviewer.allencell.org/viewer?url=https%3A%2F%2Fallencell.s3.amazonaws.com%2Faics%2Femt_timelapse_dataset%2Fdata%2F3500005829_46_raw_converted.ome.zarr&mode=maxproject&c0=cps:0:0:1:98:0:1:116:1:1:255:1:1,rmp:98:116&c1=ven:1,col:ffffff,cps:0:0:1:30:0:1:46:1:1:255:1:1,rmp:22:76&c2=col:bd10e0,cps:0:0:1:31:0:1:50:1:1:255:1:1,rmp:31:50 |
| Fig. 7A | 3D lumenoid EMT | mEGFP-tagged ZO-1 | https://volumeviewer.allencell.org/viewer?url=https%3A%2F%2Fallencell.s3.amazonaws.com%2Faics%2Femt_timelapse_dataset%2Fdata%2F3500005834_69_raw_converted.ome.zarr&mode=maxproject&c0=cps:0:0:1:84:0:1:103:1:1:255:1:1,rmp:84:103&c1=ven:1,col:ffffff,cps:0:0:1:34:0:1:45:1:1:255:1:1,rmp:23:87.279&c2=col:bd10e0,cps:0:0:1:36:0:1:59:1:1:255:1:1,rmp:36:59 |
| Fig. 7C | 2D PLF colony EMT | mEGFP-tagged ZO-1, human-specific anti- | https://volumeviewer.allencell.org/viewer?url=https%3A%2F%2Fallencell.s3.amazonaws.com%2Faics%2Femt_timelapse_dataset%2Fdata%2F3500005698_13_raw_converted.ome.zarr&mode=maxproject&reg=0:1,0:1,0.5:0.633&slice=0.5,0.5,0.567&t=34&c0=cps:0:0:1:138:0:1:151:1:1:255:1:1,rmp:138:151&c1=ven:1,col:7ed321,cps:3.1145:0:1:20.863:0:1:28.878:1:1:149.11:1:1,rmp:11.966:51.303&c2=ven:1,col:bd10e0, |

| | | | |
|---|---|---|---|
| | | collagen IV antibody | cps:-0.039842:0:1:0.89527:0:1:1.3406:1:1:2.2311:1:1,rmp:0.89527:1.5812 |
| Fig. 7C | 2D colony EMT | mEGFP-tagged ZO-1, human-specific anti-collagen IV antibody | https://volumeviewer.allencell.org/viewer?url=https%3A%2F%2Fallencell.s3.amazonaws.com%2Faics%2Femt_timelapse_dataset%2Fdata%2F3500005829_46_raw_converted.ome.zarr&mode=maxproject&reg=0:1,0:1,0.733:0.867&t=34&c0=cps:0:0:1:98:0:1:116:1:1:255:1:1,rmp:98:116&c1=ven:1,col:7ed321,cps:-0.45293:0:1:21.288:0:1:32.883:1:1:184.34:1:1,rmp:12.591:57&c2=ven:1,col:bd10e0,cps:1.3817:0:1:11.945:0:1:18.42:1:1:88.275:1:1,rmp:10.869:26.059 |
| Fig. 7C | 3D lumenoid EMT | mEGFP-tagged ZO-1, human-specific anti-collagen IV antibody | https://volumeviewer.allencell.org/viewer?url=https%3A%2F%2Fallencell.s3.amazonaws.com%2Faics%2Femt_timelapse_dataset%2Fdata%2F3500005834_69_raw_converted.ome.zarr&mode=maxproject&reg=0:1,0:1,0.46667:0.6&slice=0.5,0.5,0.53333&t=44&c0=cps:7.4059:0:1:87.963:0:1:106.18:1:1:251.95:1:1,rmp:34.87:145.64&c1=ven:1,col:7ed321,cps:4.4348:0:1:61.791:0:1:80.348:1:1:434.61:1:1,rmp:52.185:118&c2=ven:1,col:bd10e0,cps:1.1806:0:1:16.447:0:1:26.201:1:1:109.32:1:1,rmp:6:51.729 |
| Fig. 7D | 2D PLF colony EMT, 2D colony EMT, 3D lumenoid EMT | mEGFP-tagged ZO-1 | see Supplemental Movies 4-6 |
| **Figure 8: Basement membrane detachment from the glass depends on the adhesiveness of the substrate in 2D colonies and size of the lumenoid in 3D.** | | | |
| *Section* | *EMT condition* | *Structure* | *3D Volume Viewer link* |
| Fig. 8A | 2D PLF colony EMT | human-specific anti-collagen IV antibody | https://volumeviewer.allencell.org/viewer?url=https%3A%2F%2Fallencell.s3.amazonaws.com%2Faics%2Femt_timelapse_dataset%2Fdata%2F3500006212_41_raw_converted.ome.zarr&mode=maxproject&reg=0:1,0:1,0.26666666666666666:0.43333333333333335&slice=0.5,0.5,0.43333333333333335&t=24&view=Z&c0=cps:0:0:1:99:0:1:111:1:1:255:1:1,rmp:99:111&c1=col:7ed321,cps:0:0:1:42:0:1:60:1:1:255:1:1,rmp:26:71.783&c2=ven:1,col:ffffff,cps:0:0:1:19.41:0:1:33.681:1:1:98.513:1:1,rmp:2.2586:62.631 |
| Fig. 8B | 2D colony EMT | human-specific anti-collagen IV antibody | https://volumeviewer.allencell.org/viewer?url=https%3A%2F%2Fallencell.s3.amazonaws.com%2Faics%2Femt_timelapse_dataset%2Fdata%2F3500006079_10_raw_converted.ome.zarr&mode=maxproject&slice=0.5,0.5,0.4&t=24&view=Z&c0=cps:0:0:1:99:0:1:116:1:1:255:1:1,rmp:99:116&c1=col:6fba11,cps:0:0:1:50:0:1:73:1:1:255:1:1,rmp:50:73&c2 |

| | | | |
|---|---|---|---|
| | | | =ven:1,col:ffffff,cps:0:0:1:44.914:0:1:74.839:1:1:195.33:1:1,rmp:0:79.531 |
| Fig. 8C | 3D lumenoid EMT (small lumenoid) | human-specific anti-collagen IV antibody | https://volumeviewer.allencell.org/viewer?url=https%3A%2F%2Fallencell.s3.amazonaws.com%2Faics%2Femt_timelapse_dataset%2Fdata%2F3500006079_22_raw_converted.ome.zarr&mode=maxproject&slice=0.5,0.5,0.167&t=24&view=Z&c0=cps:0:0:1:100:0:1:119:1:1:255:1:1,rmp:100:119&c1=col:7ed321,cps:-10.951:0:1:75.828:0:1:108.53:1:1:309.75:1:1,rmp:75.828:108.53&c2=ven:1,col:ffffff,cps:4.1869:0:1:55.409:0:1:85.354:1:1:205.14:1:1,rmp:15.268:93 |
| Fig. 8D | 3D lumenoid EMT (large lumenoid) | human-specific anti-collagen IV antibody | https://volumeviewer.allencell.org/viewer?url=https%3A%2F%2Fallencell.s3.amazonaws.com%2Faics%2Femt_timelapse_dataset%2Fdata%2F3500006079_61_raw_converted.ome.zarr&mode=maxproject&slice=0.5,0.5,0.167&t=24&view=Z&c0=cps:-9.4873:0:1:93.866:0:1:129.87:1:1:286.64:1:1,rmp:93.866:129.87&c1=col:6fba11,cps:0:0:1:68:0:1:105:1:1:255:1:1,rmp:68:105&c2=ven:1,col:ffffff,cps:0:0:1:20.172:0:1:34.431:1:1:142.8:1:1,rmp:8.9630:73.606 |
| Fig. 8E | 3D lumenoid EMT (smallest lumenoid) | mEGFP-tagged E-cadherin, human-specific anti-collagen IV antibody | https://volumeviewer.allencell.org/viewer?url=https%3A%2F%2Fallencell.s3.amazonaws.com%2Faics%2Femt_timelapse_dataset%2Fdata%2F3500006079_27_raw_converted.ome.zarr&mode=maxproject&slice=0.5,0.45833,0.066667&cam=pos:0:2:0,up:0:0:1&view=Y&c0=cps:0:0:1:87:0:1:100:1:1:255:1:1,rmp:87:100&c1=ven:1,col:6fba11,cps:0:0:1:67:0:1:95:1:1:255:1:1,rmp:58.348:130.18&c2=ven:1,col:bd10e0,cps:0:0:1:50:0:1:77:1:1:255:1:1,rmp:32.131:98.673 |
| Fig. 8E | 3D lumenoid EMT | mEGFP-tagged E-cadherin, human-specific anti-collagen IV antibody | https://volumeviewer.allencell.org/viewer?url=https%3A%2F%2Fallencell.s3.amazonaws.com%2Faics%2Femt_timelapse_dataset%2Fdata%2F3500006079_22_raw_converted.ome.zarr&mode=maxproject&slice=0.5,0.49038,0.5&cam=pos:0:2:0,up:0:0:1&view=Y&c0=cps:0:0:1:100:0:1:119:1:1:255:1:1,rmp:100:119&c1=ven:1,col:6fba11,cps:0:0:1:69:0:1:95:1:1:255:1:1,rmp:47:132.63&c2=ven:1,col:bd10e0,cps:0:0:1:65:0:1:103:1:1:255:1:1,rmp:94.343:128.53 |
| Fig. 8E | 3D lumenoid EMT | mEGFP-tagged E-cadherin, human-specific anti-collagen IV antibody | https://volumeviewer.allencell.org/viewer?url=https%3A%2F%2Fallencell.s3.amazonaws.com%2Faics%2Femt_timelapse_dataset%2Fdata%2F3500006079_23_raw_converted.ome.zarr&mode=maxproject&slice=0.5,0.49679,0.5&cam=pos:0:2:0,up:0:0:1&view=Y&c0=cps:0:0:1:103:0:1:133:1:1:255:1:1,rmp:103:133&c1=ven:1,col:6fba11,cps:0:0:1:64:0:1:94:1:1:255:1:1,rmp:63.404:103.58&c2=ven:1,col:bd10e0,cps:0:0:1:32:0:1:52:1:1:255:1:1,rmp:32:56.743 |
| Fig. 8E | 3D lumenoid EMT | mEGFP-tagged E-cadherin, human-specific anti-collagen IV antibody | https://volumeviewer.allencell.org/viewer?url=https%3A%2F%2Fallencell.s3.amazonaws.com%2Faics%2Femt_timelapse_dataset%2Fdata%2F3500006079_25_raw_converted.ome.zarr&mode=maxproject&slice=0.5,0.52244,0.5&cam=pos:0:2:0,up:0:0:1&view=Y&c0=cps:0:0:1:84:0:1:106:1:1:255:1:1,rmp:84:106&c1=ven:1,col:6fba11,cps:0:0:1:49:0:1:81:1:1:255:1:1,rmp:58.531:78.548&c2=ven:1,col:bd10e0,cps:0:0:1:30:0:1:49:1:1:255:1:1,rmp:30:61.924 |

| | | | |
|---|---|---|---|
| Fig. 8E | 3D lumenoid EMT | mEGFP-tagged E-cadherin, human-specific anti-collagen IV antibody | https://volumeviewer.allencell.org/viewer?url=https%3A%2F%2Fallencell.s3.amazonaws.com%2Faics%2Femt_timelapse_dataset%2Fdata%2F3500006079_62_raw_converted.ome.zarr&mode=maxproject&slice=0.5,0.47115,0.5&cam=pos:0:2:0,up:0:0:1&view=Y&c0=cps:0:0:1:93:0:1:122:1:1:255:1:1,rmp:93:122&c1=ven:1,col:6fba11,cps:0:0:1:74:0:1:116:1:1:255:1:1,rmp:84.705:114.17&c2=ven:1,col:bd10e0,cps:0:0:1:45:0:1:71:1:1:255:1:1,rmp:45:71 |
| Fig. 8E | 3D lumenoid EMT | mEGFP-tagged E-cadherin, human-specific anti-collagen IV antibody | https://volumeviewer.allencell.org/viewer?url=https%3A%2F%2Fallencell.s3.amazonaws.com%2Faics%2Femt_timelapse_dataset%2Fdata%2F3500006079_61_raw_converted.ome.zarr&mode=maxproject&slice=0.5,0.50641,0.5&cam=pos:0:2:0,up:0:0:1&view=Y&c0=cps:0:0:1:89:0:1:120:1:1:255:1:1,rmp:89:120&c1=ven:1,col:6fba11,cps:0:0:1:68:0:1:105:1:1:255:1:1,rmp:84.341:100.29&c2=ven:1,col:bd10e0,cps:0:0:1:40:0:1:65:1:1:255:1:1,rmp:40:65 |
| Fig. 8E | 3D lumenoid EMT (largest lumenoid) | mEGFP-tagged E-cadherin, human-specific anti-collagen IV antibody | https://volumeviewer.allencell.org/viewer?url=https%3A%2F%2Fallencell.s3.amazonaws.com%2Faics%2Femt_timelapse_dataset%2Fdata%2F3500006079_63_raw_converted.ome.zarr&mode=maxproject&slice=0.5,0.52244,0.5&cam=pos:0:2:0,up:0:0:1&view=Y&c0=cps:0:0:1:89:0:1:123:1:1:255:1:1,rmp:89:123&c1=ven:1,col:6fba11,cps:5.8621:0:1:35.172:0:1:53.931:1:1:105.52:1:1,rmp:40.209:58.97&c2=ven:1,col:bd10e0,cps:-0.62808:0:1:38.404:0:1:77.436:1:1:765:1:1,rmp:38.404:73.494 |
